# Seed or soil: tracing back the plant mycobiota primary sources

**DOI:** 10.1101/2023.08.04.551828

**Authors:** Liam Laurent-Webb, Kenji Maurice, Benoît Perez-Lamarque, Amélia Bourceret, Marc Ducousso, Marc-André Selosse

## Abstract

Plants host diverse communities of fungi (collectively called the mycobiota) which play crucial roles in their development. The assembly processes of the mycobiota, however, remain poorly understood, in particular, whether it is transmitted by parents through the seeds (vertical transmission) or recruited in the environment (horizontal transmission). Here we attempt to quantify the relative contributions of horizontal and vertical transmission in the mycobiota assembly of a desert shrub, *Haloxylon salicornicum,* by comparing the mycobiota of *in situ* bulk soil and seeds to that of (i) *in situ* adult individuals and (ii) *in vitro*-germinated seedlings in soil collected *in situ*, either autoclaved or not. We show that the mycobiota is partially transmitted through the seeds to seedlings. In contrast, root mycobiota of adults are highly similar to that of bulk soil, whereas adult leaf mycobiota remain similar to that of seeds. Thus, the mycobiota is transmitted both horizontally and vertically depending on the plant tissue. Despite discrepancies between *in situ* and *in vitro* approaches, our result may also suggest a compositional turnover in plant mycobiota during plant development. Understanding the respective contribution of these transmission paths to the plant mycobiota is fundamental to deciphering potential coevolutionary processes between plants and fungi.

## Introduction

All plants are colonized on their surface and in their tissues by diverse communities of microorganisms, such as bacteria (Bulgarelli *et al*., 2013) and fungi (Selosse *et al*., 2004; Rodriguez *et al*., 2009). Collectively, these microorganisms are referred to as the microbiota (Berg *et al*., 2020) for which plants tend to have developed functional dependency (Selosse *et al*., 2014). Among the plant microbiota, fungi form the mycobiota which play key roles in the plant life cycle as they interact with their plant host in various manners, ranging from parasitic to mutualistic (Trivedi *et al*., 2020). Though fungi are well-known to have deleterious effects on plant fitness (e.g., pathogenic fungi), it has been recognized over the past decades that they are also involved in many crucial plant functions such as nutrition (Smith and Read, 2009; Yakti *et al*., 2018), defense against pathogens (Rodriguez Estrada *et al*., 2012), resistance to various stresses (Li *et al*., 2018; Hosseyni *et al*., 2021), and ecological success (Selosse *et al*., 2014). Some fungi form specialized structures with their host plant such as mycorrhizae (van der Heijden *et al*., 2015), but others colonize their hosts with no apparent symptoms, such as endophytic fungi (*sensu* Wilson, 1995). Plants and their microbiota are sometimes referred to as ‘holobionts’ (e.g., Vandenkoornhuyse *et al*., 2015), a theoretical framework stipulating that the accumulation of the plant organism and its microbiota (holobiont) form a unit of selection. This notion is still subject to debate as it implies fidelity between the partners; yet, mutualistic associations tend to be generalist and in most cases are not vertically transmitted across generations (Bright and Bulgheresi, 2010; Douglas and Werren, 2016).

Partners’ fidelity may be guaranteed by vertical transmission of the mycobiota (Wilkinson, 1997), i.e., if the plant-associated fungi are transmitted from generation to generation by way of seeds (Bright and Bulgheresi, 2010) and/or during vegetative multiplication (Vannier *et al*., 2018). Thanks to novel DNA sequencing and barcoding technologies, healthy seeds are no longer considered sterile, as they generally present a diverse and dynamic microbiota at their different developmental stages (Klaedtke *et al*., 2016; Nelson, 2018; Abdelfattah *et al*., 2022; Simonin *et al*., 2022), so that fungal vertical transmission through the seeds is possible. However, the sources and mechanisms of acquisition of plant-associated fungi are still poorly understood despite a growing interest in these organisms (Harrison and Griffin, 2020). Recent findings suggest that fungi may be acquired by the plant (i) directly from their environment (horizontal transmission, from the soil for instance; Bonito *et al*., 2014) or (ii) from their parents (vertical transmission) through the seeds (Gundel *et al*., 2011; Shade *et al*., 2017) or clonal structures (e.g., aerial stolons; Vannier *et al*., 2018). In particular, most fungi colonizing belowground tissues (rhizosphere, roots) are considered to be mainly horizontally transmitted from the soil (Lundberg *et al*., 2012; Bonito *et al*., 2014). Yet, root-associated fungi may also be indirectly transmitted to seedlings by secondarily colonizing fruit or seed tissues, such as *Scleroderma bermudense* spores carried by *Cocoloba uvifera* dried fruits (Séne *et al*., 2018), a process referred to as pseudo-vertical transmission. Cases of direct vertical transmission have also been demonstrated for several fungal leaf endophytes such as grass endophytes in the genus *Epichloe* (Tintjer *et al*., 2008; Rodriguez *et al*., 2009) or *Alternaria alternata* and *Cladosporium sphaerospermum* in several plant species (Hodgson *et al*., 2014), suggesting a common phenomenon in aerial tissues (see Class 1 and Class 2 endophytes as defined by Rodriguez *et al*., 2009). Vertical transmission was also confirmed by *in vitro* experiments demonstrating that both bacterial and fungal microbiota may be partially transmitted from seeds to seedlings (Johnston-Monje *et al*. (2016) in maize rhizosphere; Robinson *et al*. (2016) in wheat roots; Abdelfattah *et al*. (2021) in above- and belowground oak tissues; Barret *et al*. (2015) in several Brassicaceae species, Rezki *et al*. (2018) in radish seeds over several generations), sometimes over several plant generations (e.g., in above- and belowground rice tissues (Hardoim *et al*., 2012) or in *Setaria veridis* seeds (Rodríguez *et al*., 2020).

Available knowledge therefore suggests that a seedling’s mycobiota originate from both seeds and soil in proportions that vary between aerial and belowground tissues. The respective contribution of these two sources to the microbiota are, however, often considered separately. For instance, Xiong *et al*. (2021) studied the contribution of soil to the bacterial microbiota of maize (*Zea mays*), wheat (*Triticum aestivum*), and barley (*Hordeum vulgare*) and showed that soil (horizontal transmission) is the primary reservoir of the bacterial microbiota even in aerial compartments. Only a few studies have comparatively qualified and quantified the vertical and horizontal transmission of the microbiota (see Moroenyane *et al*., 2021; Walsh *et al*., 2021 for bacteria), especially in fungi (but see Rochefort *et al*., 2021). These studies highlight that seedling bacterial microbiota are mainly influenced by seeds (Moroenyane *et al*., 2021 in soybean; Walsh *et al*., 2021 in wheat), while the soil is the main source of seedling mycobiota (Rochefort *et al*., 2021 in *Brassica napus*). However, these studies only used *in vitro* experimental designs, which may not reflect *in situ* conditions (e.g., complex plant communities, differences in microbiota composition between substrates used *in vitro* and soil *in situ*…) and therefore influence transmission pathways.

Desert ecosystems represent one-third of the world’s land surface (Prăvălie, 2016), and their area is expected to increase under climate change (Intergovernmental Panel on Climate Change, 2022). They are characterized by low precipitation and nutrient availability resulting in lower fungal (Tedersoo *et al*., 2014) and plant (Cai *et al*., 2023) diversity compared to other biomes. In hot deserts, vegetation is often discontinuous and patchy, with perennial species regularly spaced and separated by bare soil (sometimes colonized by annual plants; de Graaff *et al*., 2014). This patchy distribution of shrubs is referred to as ‘fertility islands’ or ‘resource islands’: soil nutrient concentrations (i.e., C, N, P) close to shrubs are higher than in surrounding bare soils (Schlesinger and Pilmanis, 1998). These fertility islands are therefore hotspots of microbial diversity, especially for fungi (Ochoa-Hueso *et al*., 2018; Maurice *et al*., 2023). As they display rather simple and spatially structured plant communities, desert ecosystems are interesting models for *in situ* ecological research. The patchy distribution of shrubs may limit the impact of other neighboring plant species on their mycobiota and in particular the influence of conspecific individuals (Schneider-Maunoury *et al*., 2020; Brigham *et al*., 2023). Moreover, the harsh soil conditions may limit microbial availability in the environment and favor vertical transmission.

Here, we take advantage of a desert ecosystem to quantify both vertical and horizontal transmission pathways during mycobiota assembly. We hypothesize that (i) plant mycobiota are mainly influenced by the soil mycobiota with a low contribution of seeds, even at early development stages, and that (ii) different compartments (especially aerial and underground ones) show contrasted patterns of colonization from seeds and soil. We studied *Haloxylon salicornicum* (Amaranthaceae) as a model, a common desert shrub of the Eastern Arabian flora (Al Salameen *et al*., 2018) with potential for restoration and protection of arid lands (Rathore *et al*., 2015). We assessed the contribution of both horizontal (soil) and vertical (seeds) transmission in the mycobiota assembly of *Haloxylon salicornicum* through the comparison of mycobiota compositions obtained (i) by sampling bulk soil *in situ* and the different compartments of adult individuals *in situ* (rhizosphere, roots, leaves, and seeds) and (ii) by germinating seeds *in vitro* in non-processed and autoclaved bulk soil collected *in situ*.

## Material & Methods

### *In situ* sampling

Bulk soil (i.e., soil without apparent vegetation), rhizosphere, roots, and leaves of *in situ Haloxylon salicornicum* samples were collected at 5 sites in the Shaaran Natural Reserve (Province of Medina, AlUla, Saudi Arabia; Fig. 1). Bulk soil, rhizosphere, and root samples were collected in March 2022 (Supp. Table 1). At each of the 5 sites, 5 bulk soil samples and 13 roots and rhizospheric soil (i.e., soil attached and/or closed to the roots) were collected (Table 1; Fig. 1). Roots and rhizosphere samples were collected from the same individual each time (Fig. 1). Soil samples (bulk and rhizosphere) were sieved to 2 mm and root samples were stored in a 2% CTAB solution. All samples were stored at 4°C until molecular analyses. In May 2022, we collected samples of the aerial parts of the same individuals than in March 2022. As *H. salicornicum* harbors small reduced leaves on green photosynthetic stems (Singh *et al*., 2015), we sampled both stems and leaves together and will refer to them as *leaves*. Sampled leaves were dried using silica gel and kept dry until molecular analyses. We studied both epiphytic and endophytic leaf mycobiota. We also collected additional bulk soil from two natural sites (that is, sites with no apparent influence of human activities) for the *in vitro* experiment (3 bulk soil samples per site, pooled). Bulk soil samples collected for the *in vitro* experiment were dry when sampled (data not shown). They were unprocessed and kept dry for one week before setting up the *in vitro* experiment. Seeds were collected during spring 2022 on *H. salicornicum* individuals from previously studied sites before contact with soil. We randomly sampled 20 pools of 9 seeds from this seed collection for molecular analysis and randomly sampled ca. 200 additional seeds for *in vitro* experiments.

**Figure 1:**
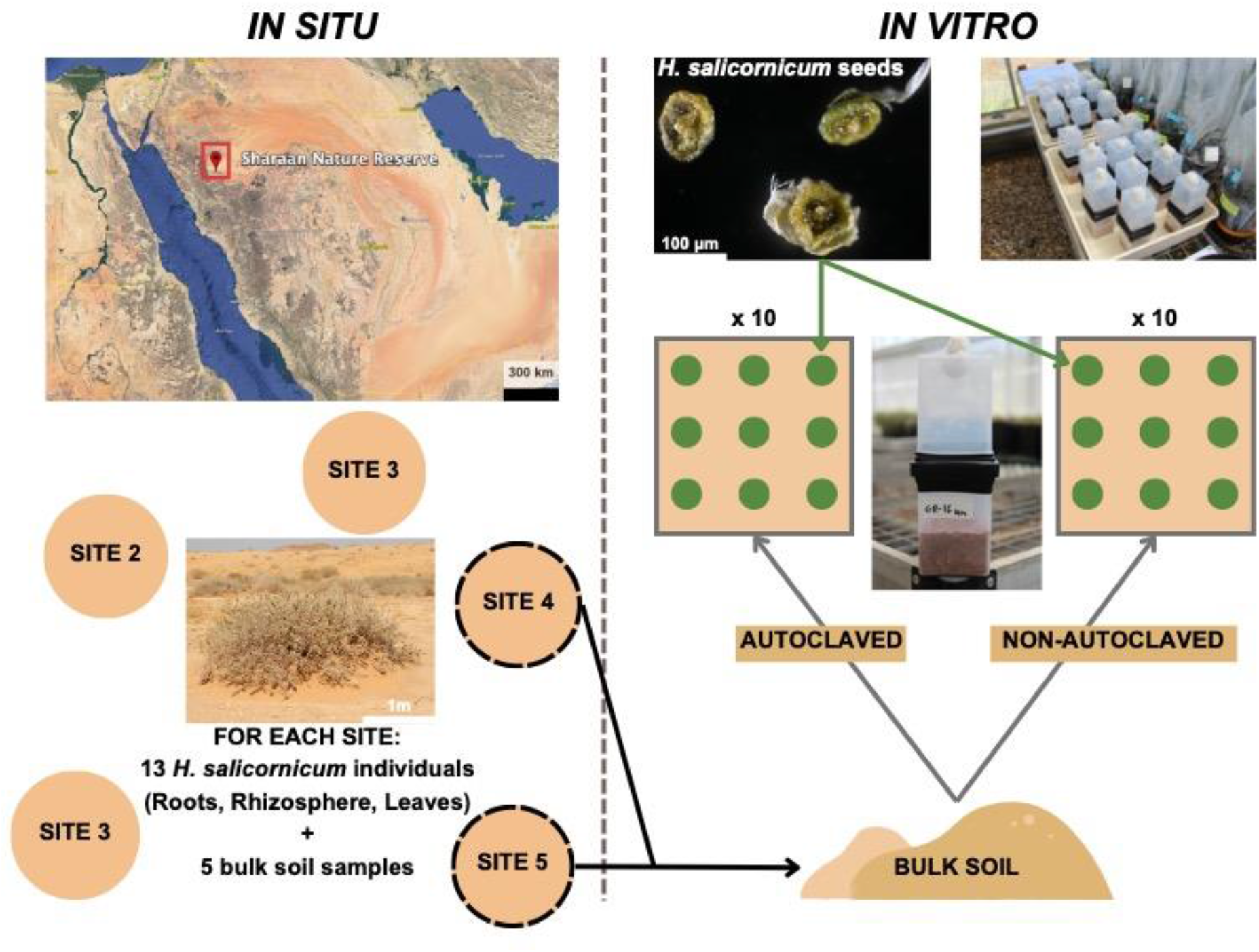
Experimental design of the study. Samples were collected at five sites in the Shaaran Nature Reserve, Province of Medina, Saudi Arabia. At each site, rhizosphere, roots, and leaves of 13 *H. salicornicum* individuals were sampled along with five samples of bulk soil (i.e., soil without apparent vegetation). Additional bulk soil was collected at two of the five sites *in situ* and pooled. We germinated seeds in a heated glass house on the additional bulk soil collected *in situ,* either autoclaved or not.

**Table 1:** Soil and *H. salicornicum* samples collected.

|  |  | Bulk soil | Rhizosphere | Roots | Leaves | Seeds |
| --- | --- | --- | --- | --- | --- | --- |
| Collected samples | *In situ* | 25 | 65 | 65 | 65 | 20 (pools of 9 seeds) |
|  | *In vitro* |  |  | 20<br>(13 + 7) | 20<br>(13 + 7) |  |

### *In vitro* germination experiment

We germinated seeds in a heated glass house with a temperature of 25°C-28°C, low humidity, and day/night alternations of respectively 16 h and 8 h. We used bulk soil collected *in situ* as substrate (Fig. 1). Bulk soil samples collected for the *in vitro* experiment were pooled (see *In* situ sampling) and kept unprocessed for one week before setting up the experiment. Pooled bulk soils were either autoclaved (30 min at 120°C) or not, leading to the two conditions of the *in vitro* experiment (resp. autoclaved and non-autoclaved). We expect this autoclaved condition to eliminate most of the fungi present in bulk soil (although some might resist; Wolf *et al*., 1989). For each condition, 10 Magenta boxes (Merck, Germany) were filled with 250g of bulk soil as substrate. Before planting, we removed petals and sepals from seeds with sterile tweezers. Nine seeds per Magenta box were planted at equal distance using sterile tweezers (Fig. 1). Substrate was irrigated with sterile Milli-Q water (40 mL/box). Boxes were closed with another Magenta box, with a 1 cm diameter hole filled with cotton to allow gas exchange and limit aerial contaminants. Of the 180 seeds initially planted, 13 reached the 2-leaf stage in the autoclaved condition and 7 in the non-autoclaved condition (Table 1; differences in germination rate were not significant, χ^2^ test: *p* = 0.24). As the germination rate was low and because we experienced some damping-off at a later stage during previous trials (data not shown), leaves and roots of seedlings were collected after 7 days (2-leaf growth stage) under sterile conditions. Neither roots nor leaves showed signs of infection or disease. Roots were washed with sterile water and all samples (roots and leaves) were flash-frozen with liquid nitrogen. Samples were kept at −20°C until molecular analyses.

For *in vitro* samples, the left number in brackets is the number of samples in the autoclaved condition, and the right number (underlined) is the number of samples in the non-autoclaved condition.

### Molecular analyses

*In situ* roots were washed with sterile water, flash-frozen with liquid nitrogen and ground using a ceramic ball for 3 x 30s at speed 5.0 in a FasPrep-24^TM^ 5G instrument (MP Biomedicals^TM^, Solon, USA). Leaves (*in situ* and *in vitro*), roots (*in vitro*), and seeds (*in situ*) were ground using two sterile stainless-steel beads in a Tissue Lyzer II (Qiagen, Germany) for 3×30s at 30Hz. To increase DNA concentration, 5 mL of soil was mixed with 20 mL of sterile Milli-Q water. After 1h of mixing, the supernatant was isolated and centrifuged at 12,000 g. The resulting pellet was isolated for extraction. DNA extraction of all samples was performed with the FastDNA Spin Kit for soil (MP Biomedicals^TM^, Solon, USA) following the manufacturer’s instructions. DNA concentration of extracts was measured with the PicoGreen fluorophore (Quant-iT™ PicoGreen™ dsDNA Assay Kits, Thermo Fisher Scientific, USA). Bulk soil samples, roots, and rhizosphere had low DNA concentrations and were therefore normalized to 0.3 ng.µL^-1^. Seed and leaf extracts showed higher DNA concentrations (up to 150 ng.µL^-1^; data not shown). We therefore diluted these extracts to 3.5 ng.µL^-1^. The ITS2 region of the fungal ribosomal operon was amplified for all samples with tagged primers ITS86F (5′-GTGAATCATCGAATCTTTGAA-3′) and ITS4 (5′-TCCTCCGCTTATTGATATGC-3′; White *et al*., 1990; Op De Beeck *et al*., 2014). Reactions were performed using the Thermo Scientific Phusion™ High-Fidelity DNA Polymerase (Thermo Fisher Scientific, USA). All reactions were performed in pseudo-triplicate. Triplicates were then pooled and checked on 2% agarose gel. PCR products were purified using Agencourt AMPure XP beads (Beckman Coulter Inc., Indianapolis, IN, USA) and their concentrations were measured with PicoGreen. We built two equimolar pools for sequencing: (1) PCR products from *in situ* samples of bulk soil, rhizosphere, and roots were mixed in one equimolar pool whereas (2) samples of leaves *in situ* and *in vitro*, roots *in vitro*, and seeds were pooled in another equimolar pool. The two pools were purified twice with AMPure. Each pool was sequenced independently using *MetaFast* library preparation and sequencing (performed by Fasteris SA, Switzerland) on an Illumina platform using the 2×250 pb Miseq technology (Illumina).

### Bioinformatic analysis

A pipeline based on VSEARCH (Rognes *et al*., 2016) and available on GitHub (https://github.com/BPerezLamarque/Scripts/) was used for data processing. Briefly, paired-end reads were merged and quality checked. Merged reads were then demultiplexed using *cutadapt* (Martin, 2011) with 0 error accepted in primer or tag sequences. Reads from all samples were dereplicated and clustered as classical 97% sequence similarity Operational Taxonomic Units (OTUs) using VSEARCH as recommended in Tedersoo *et al*. (2022). All sequences were checked for the presence of chimeras. The taxonomy of these OTUs was assigned with VSEARCH against the UNITE v9.0 database (Nilsson *et al*., 2019). Reads were filtered to keep only non-chimeric sequences of > 200 pb and with a total abundance of at least 10 (see Supp. Methods 2 for details on the functions and parameters used). We used the *decontam* algorithm (Susana Rivera *et al*., 2011) to remove potential contaminants in the datasets, using both *prevalence* and *frequency* algorithms (Supp. Methods 2). Samples with less than 1,000 fungal reads were discarded. After filtering, we obtained the mycobiota composition of 259 samples (96% of all collected samples; Supp. Table 1), with a mean sequencing depth of 21,013 reads per sample (ranging from 1,041 to 130,227; Supp. Fig. 1 & 2). In order to compute UniFrac distances, we reconstructed the fungal phylogenetic trees as in Perez-Lamarque *et al*. (2022) (Supp. Methods 2).

### Statistical analysis

OTU tables were processed using the *phyloseq* package (McMurdie and Holmes, 2013) in R (R Core Team, 2023).

#### Richness and diversity analysis

We computed richness (Chao1 estimator) and diversity (Shannon index) based on actual counts using the *vegan* R package (Oksanen *et al*., 2013). In order to test differences between experimental designs (*in situ* and *in vitro*), compartments (bulk soil, rhizosphere, roots, leaves, and seeds) and substrate conditions *in vitro* (non- and autoclaved bulk soil), we used linear regression and Tukey’s post hoc pairwise test (Supp. Methods 3).

#### Assessing differences in community structure

Beta-diversity analyses were performed using either relative abundances (as they may perform better for community comparisons; Gloor *et al*., 2017; McKnight *et al*., 2019) or the Hellinger-transformed data to correct for variability in sampling depth (Legendre and Gallagher, 2001). We computed Bray-Curtis distances using the *vegan* package (Oksanen *et al*., 2013). We also computed UniFrac distances in order to account for the phylogenetic relatedness between OTUs in our community composition and clustering analyses. We visualized community composition differences using principal component analysis (PCoA). As both transformations (relative abundance and Hellinger transformation) and both distances (Bray-Curtis and UniFrac) showed similar results, we only report results from Bray-Curtis distances from relative abundances in the main text. In order to test for differences in mycobiota composition between the two experimental designs and between compartments, we used *PERMANOVA* (10,000 permutations) with all samples using the following model: *distance*∼*exp. design*\**compartment* (where *exp. design* corresponds to *in situ* or *in vitro*). Then, to test whether the two substrate conditions led to different mycobiota composition and to test for differences in composition between above- and belowground compartments, we used a second *PERMANOVA* (10,000 permutations) with only leaves and roots of individuals germinated *in vitro* using the following model: *distance*∼*compartment*\**substrate condition*.

#### Construction of plant/fungal bipartite networks to assess the sharing of OTUs

To further assess the sharing of OTUs between samples (in particular between bulk soil, seeds, and other plant tissues), we constructed bipartite networks using the *igraph* R package (Csardi and Nepusz, 2006). Nodes represent OTUs and samples, and the edge width indicates the relative abundance of an OTU in a sample. The size of the nodes is proportional to their betweenness centrality, a measure of centrality based on shortest paths between nodes. Bipartite networks are visualized using the Fruchterman-Reingold algorithm (Fruchterman and Reingold, 1991). In particular, we want to assess whether plant/fungus networks are specialized (i.e., whether samples from the same compartment tend to share more OTUs than expected by chance, if OTUs are arbitrarily assigned to the samples) or not and which compartments may share more fungal OTUs. If horizontal transmission is the main transmission pathway, we expect that bulk soil samples and plant tissues share a lot of OTUs, while seeds and plant tissues would share more OTUs if vertical transmission dominates. We built one network with all samples and a subnetwork to give a clearer representation of the *in vitro* experiment (with seedlings, seeds, and bulk soils used for this experiment). In order to test whether OTUs are shared preferentially between samples from the same compartments or not, we built one network for each compartment of the *in situ* dataset and one network by compartment x substrate condition combinations in the *in vitro* experiment to allow for comparisons of specialization between compartments and substrate conditions. For each sample, we consider an interaction with a fungal OTU as long as the latter represents at least 0.5% of the reads. We computed the connectance C (i.e., the proportion of realized links) and the network specialization index H_2_’ (degree of specialization of the entire network) of each network. Connectance ranges from 0 to 1; values close to 1 indicate that a high proportion of all potential links are realized. H_2_’ also ranges from 0 to 1, 0 indicating a strong generalization and 1 that the network is globally specialized. H_2_’ is robust against differences in sampling size and is therefore suited for network comparisons (Blüthgen *et al*., 2006). Significance of both connectance (C) and specialization (H_2_’) was tested using null models (Supp. Methods 3).

#### Source tracking analysis

To estimate the respective contributions of seeds (vertical transmission) and bulk soil (horizontal transmission) to the *H. salicornicum* mycobiota, we used the *fast expectation-maximization for microbial source tracking* (FEAST) algorithm developed by Shenhav *et al*. (2019) and implemented in R. This algorithm estimates the fraction of a microbial community (the ‘sink’) that can be explain by different potential microbial sources (the ‘sources’). The algorithm also reports an unexplained fraction referred to as the ‘unknown’ source. Here, seeds and bulk soil samples were defined as ‘sources’ whereas roots, rhizosphere, and leaves were defined as ‘sinks’. We ran the procedure twice: first, on the *in situ* adult individuals and, a second time on seedlings from the germination experiment (with *in situ* seeds and bulk soil samples used for the germination experiment). We tested using linear regressions whether different sinks, experimental designs, compartments and/or substrate conditions were associated with significant differences in sources’ contributions to the mycobiota. (Supp. Methods 3).

#### Identifying potentially transmitted OTUs

We identified OTUs potentially transmitted from the sources (bulk soil and seeds) to the sinks, using a ‘strict’ and a ‘loose’ definition. With our strict definition, a potentially transmitted OTU is an OTU shared between one source and one sink (e.g., between bulk soil and rhizosphere), but absent from the other source (seeds in this example). This definition ensures that the OTUs identified are strictly transmitted from the studied source. With our loose definition, potentially transmitted OTUs are OTUs shared between one source and one sink (e.g., bulk soil and rhizosphere) but not necessarily absent from the other source (here seeds). This second definition allows us to identify ubiquitous OTUs that may be transmitted both horizontally and vertically. We then computed the mycobiota composition of each compartment in each experimental design and substrate condition (for *in vitro* samples) taking only potentially transmitted OTUs into account, and the mean share of the mycobiota they represent.

## Results

We successfully sequenced the fungal ITS2 region of 259 samples, including roots, rhizosphere, and leaves of *H. salicornicum* individuals *in situ* and their associated bulk soil (i.e., with no apparent vegetation). We also characterized the mycobiota of seeds collected in the same area and leaves and roots of *H. salicornicum* seedlings germinated *in vitro* in non- and autoclaved soil (see Supp. Table 1 for details on successfully processed samples). The mean sequencing depth for these samples was 21,013 reads (ranging from 1,041 to 130,227; Supp. Figure 1 & 2) after removing extraction and PCR contaminants. Rarefaction curves tended to reach a plateau for most of the samples (Supp. Fig. 2), suggesting that our sampling properly describes the mycobiota diversity of soil and *H. salicornicum* tissues.

**Figure 2:**
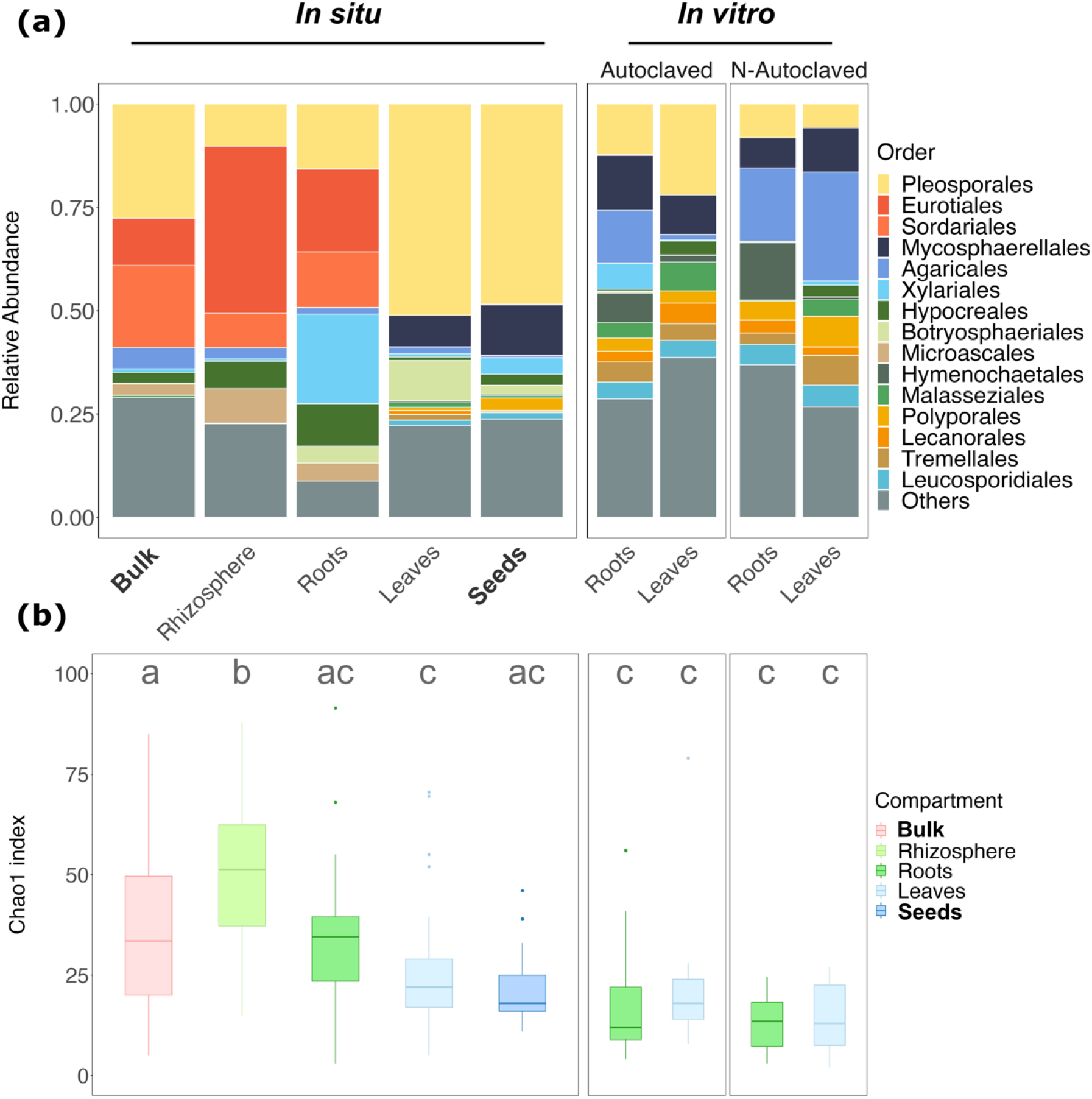
*H. salicornicum* mycobiota varies according to the compartment *in situ* but not between compartments or substrate condition *in vitro*. (a) Mycobiota composition (relative abundance, in proportions) of *H. salicornicum* compartments *in situ* and *in vitro* at the order level. Barplots represent the mean proportion of each fungal order in each compartment, experimental design, and substrate condition *in vitro*. For *in vitro* samples, both substrate conditions (non- and autoclaved) are shown. For readability purposes, only the main 15 orders are shown, while the rest are aggregated in the ‘Others’ category. **(b)** Fungal richness (Chao1 index) in the different compartments, experimental designs (*in situ* and *in vitro*) and substrate conditions *in vitro*. Different letters indicate significant differences (*p*<0.05; Tukey’s post hoc pairwise test). Sources (bulk soil and seeds) are in bold font.

### *H. salicornicum* displays contrasted mycobiota diversity between compartments in adults *in situ* but not in seedlings *in vitro*

Adult individuals *in situ* harbored contrasted mycobiota compositions depending on the compartment studied, in particular between above- and belowground compartments, whereas seedlings *in vitro* displayed similar leaf and root mycobiota in the two substrate conditions (non- and autoclaved; Fig. 2).

*In situ*, soil compartments (bulk, rhizosphere, and roots) were characterized by a greater proportion of Eurotiales and Sordariales compared to aerial compartments (leaves and seeds) which were mainly colonized by Pleosporales (representing almost 50% of the mycobiota for both aerial compartments) and to a smaller extent by Mycosphaerellales (Fig 2a). Roots were also characterized by a larger share of Xylariales (Fig. 2a) compared to other compartments. The share of Botryosphaeriales was higher in leaves than in seeds. *In vitro*, we observed small differences between seedling roots and leaves compared to the differences observed between compartments of adults *in situ*, or between substrate conditions (Fig 2a). Seedlings *in vitro* harbored a small share of Pleosporales compared to *in situ* leaves and seeds but similar proportions of Mycosphaerellales. Contrary to roots *in situ*, roots of seedlings *in vitro* harbored almost no Eurotiales or Sordariales and a small share of Xylariales. Interestingly, despite similar mycobiota composition at the order level, the top 5 OTUs were quite different between compartments and substrate conditions *in vitro*, except for *Mycosphaerella asteroma* which was ubiquitous (Supp. File 1).

The highest fungal richness was observed in the rhizosphere of adults *in situ* (mean *chao1* = 49±17; Fig. 2b), followed by bulk soil and roots *in situ* (37±22 and 33±15, respectively). Among samples *in situ*, seedling leaves and seeds harbored the lowest richness (25±13 and 22±11, respectively. All samples *in vitro* had lower richness compared to samples *in situ* (ranging from 13±8.1 for roots in the non-autoclaved condition to 22±18 for leaves in the autoclaved condition; ANOVA: *p*<10^-15^; Supp. Table 2). Differences in richness were not significant between substrate conditions *in vitro*. Similarly, differences in richness between roots and leaves *in vitro* were not significant (*p*=0.54, ANOVA; Supp. Table 2), though leaves tended to harbor a slightly greater richness (Fig. 2b; Supp. Table 2). The Shannon diversity index revealed similar trends (Supp. Fig 3; Supp. Table 3). Diversity and richness therefore displayed similar patterns as in seedlings *in vitro*; no significant differences between roots and leaves and substrate conditions were identified (Supp. Fig. 2; Supp. Table 2; Supp. Table 3), while in adults *in situ*, compartments differed significantly in richness and diversity.

**Figure 3:**
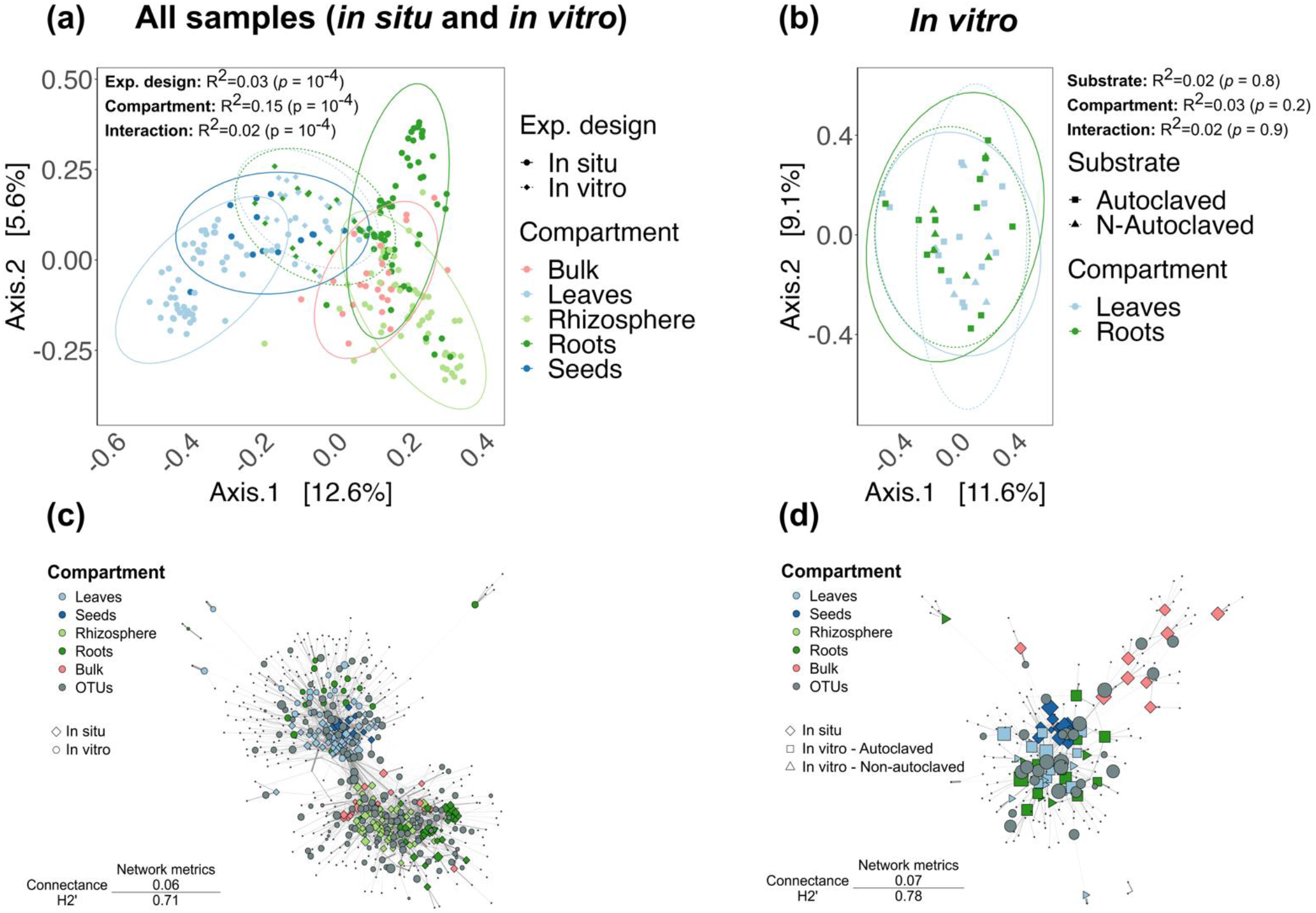
*H. salicornicum* fungal communities cluster in two main groups: soil compartments *in situ* (bulk soil, rhizosphere, roots) and aerial compartments *in situ* (leaves and seeds) plus samples *in vitro*. (a), (b) Principal coordinate analysis (PCoA) of the Bray-Curtis distances between samples. Ellipses represent the normal-probability contours of the data. Influence of the variables on Bray-Curtis distance matrices was tested using *PERMANOVA* (10 000 permutations). The procedure was run for **(a)** all samples (full lines represent *in situ* samples and dashed lines *in vitro* samples) and **(b)** samples *in vitro* only (full lines represent samples from the autoclaved substrate condition and dashed lines the non-autoclaved substrate condition). **(c), (d)** Weighted bipartite networks (using relative abundances of OTUs), visualized using the Fruchterman–Reingold layout algorithm for better readability. Grey nodes represent OTUs and colored ones represent samples. Node diameter is proportional to the betweenness centrality. Width of the edges is proportional to the relative abundance of an OTU in a sample. See Supp. Methods 3 for connectance (C) and specialization (H_2_’) calculation and significance tests. **(c)** Total dataset: round colored nodes represent samples *in vitro*, colored rhombuses represent samples *in situ*. **(d)** Samples *in vitro* with potential sources of mycobiota (bulk soil and seeds used for the experiment). Colored rhombuses correspond to *in situ* samples, squares to sample from the autoclaved condition and triangles to non-autoclaved condition. Node and edge sizes have been increased for readability purposes.

#### Mycobiota communities are highly structured in two main groups

As illustrated by the principal coordinate analysis (PCoA; Fig. 3a), compartments had distinct fungal communities (*PERMANOVA*: R^2^=0.15, *p*<10^-4^; Fig. 3a) as samples from the same compartment tended to cluster together. Differences between *in situ* and *in vitro* communities were also observed, as leaves and roots *in vitro* did not cluster with leaves and roots *in situ* (R^2^=0.03, *p*<10^-4^; Fig 3a). Globally, *in situ* soil compartments tended to cluster together, while *in situ* aerial compartments and *in vitro* samples tended to cluster in a second group (Fig. 3a). We ran the analysis with only *in vitro* samples in order to test for differences between compartments and substrate conditions (non- and autoclaved soil) and we observed no significant influence of the compartment nor substrate condition (*p*>0.1 for both variables and their interaction term; Fig. 3b). These results were consistent when using Hellinger-transformed data instead of relative abundances and when using UniFrac distances instead of Bray-Curtis distances (Supp. Fig. 4).

**Figure 4:**
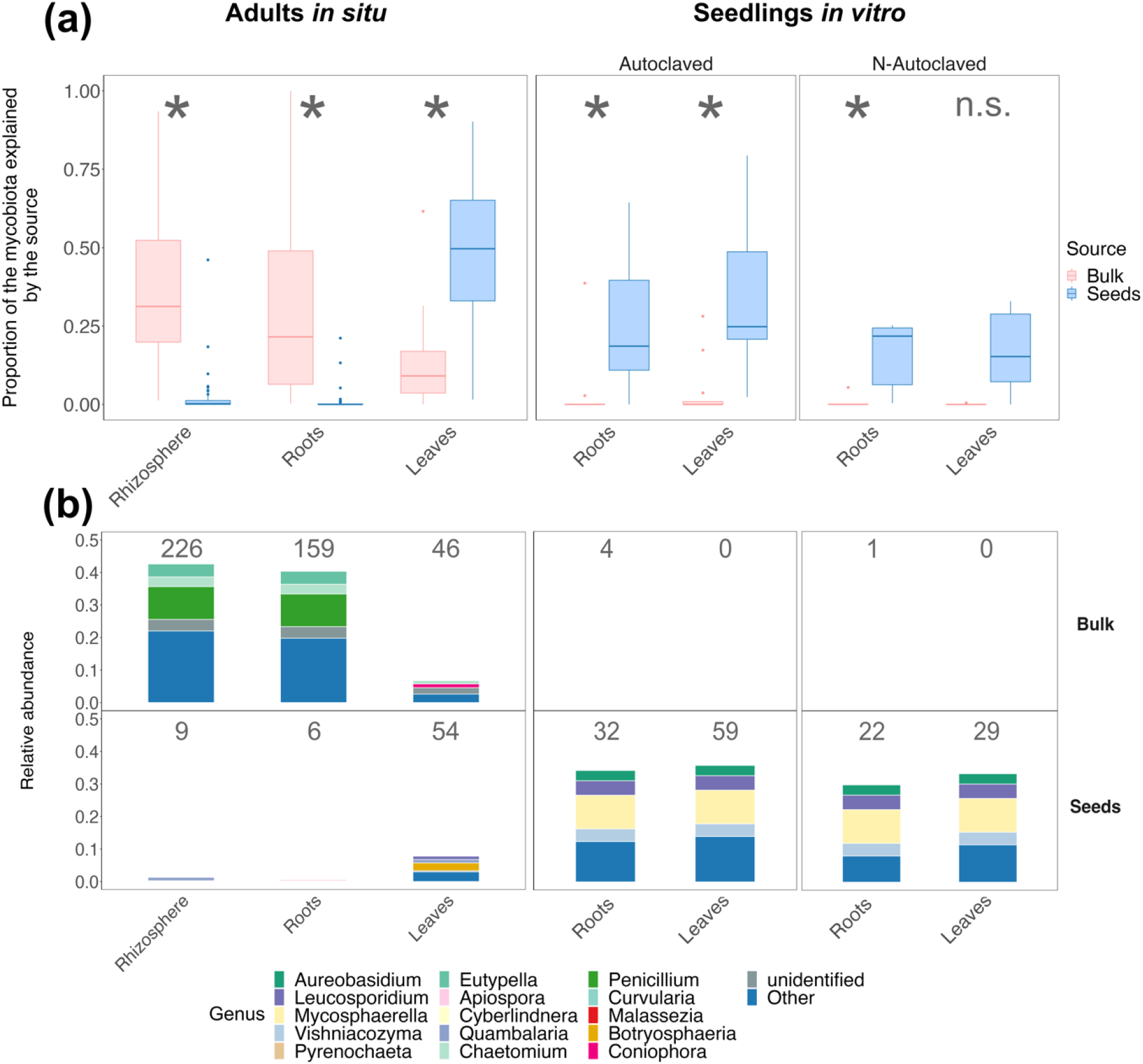
Vertical and horizontal transmissions differ between adults *in situ* and seedlings *in vitro* and between compartments, transmitting different fungal communities. **(a)** Proportion of the mycobiota of each compartment in each experimental design and substrate condition *in vitro* explained by the different potential sources (bulk soil and seeds). Proportions were estimated using the *FEAST* algorithm. Asterisks indicate significant differences between the proportions of the two sources, based on Tukey’s post hoc pairwise test (see Supp. Methods 3 & Supp. Fig. 6). **(b)** Mycobiota composition at the genus level of each compartment in each experimental design and substrate condition *in vitro* using the strict definition of potentially transmitted OTUs (i.e., we only took into consideration OTUs exclusively shared between the studied compartment and the potential source). The number of shared OTUs is indicated at the top of each bar stack.

We also constructed two bipartite networks to compare the structure of the fungal communities and assess the sharing of OTUs between compartments, experimental design and substrate condition. When considering the total network (Fig. 3c), we identified two main modules: one with *in situ* soil compartments (bulk soil, rhizosphere, roots) and another one with samples *in vitro*, seeds, and leaves *in situ* (Fig. 3c). These two modules shared some OTUs with high betweenness centrality, that is central OTUs that link many nodes of the network. The total network had a low connectance (C=0.06), meaning that a low proportion (6%) of all possible interactions between OTUs and samples was observed, which is significantly lower than expected based on our null model (*p*<0.025). Furthermore, the specialization of the network was significantly higher than expected by chance (H_2_’=0.71; *p*<0.025). Altogether, these results suggest a limited sharing of OTUs between samples. When focusing on the network formed by seedlings *in vitro*, seeds and bulk soils used as substrate for the *in vitro* experiment, seeds, and seedling tissues (leaves and roots) tended to cluster together while bulk soil samples seemed more peripheral (Fig. 3d). Samples from the non- and autoclaved conditions did not seem to form specific groups (except one outlier in the non-autoclaved condition), a result consistent with the *PERMANOVA* analysis (R^2^=0.02, *p*=0.8; Fig. 3b). When considering each compartment separately (in each substrate condition for seedlings *in vitro*), we observed that the connectance of the networks was ca. 4 times higher compared to the total network (Supp. Fig. 5), suggesting that the low connectance of the networks in Fig. 3c,d is linked to a low sharing of OTUs between compartments despite high sharing of OTUs within compartments. *In situ*, soil compartment networks tended to be more specialized (bulk: H_2_’=0.7; rhizosphere: H_2_’=0.61; roots: H_2_’=0.75; Supp. Fig. 5a) compared to aerial compartments (leaves: H_2_’=0.46; seeds: H_2_’=0.58; Supp. Fig. 5a).

**Figure 5:**
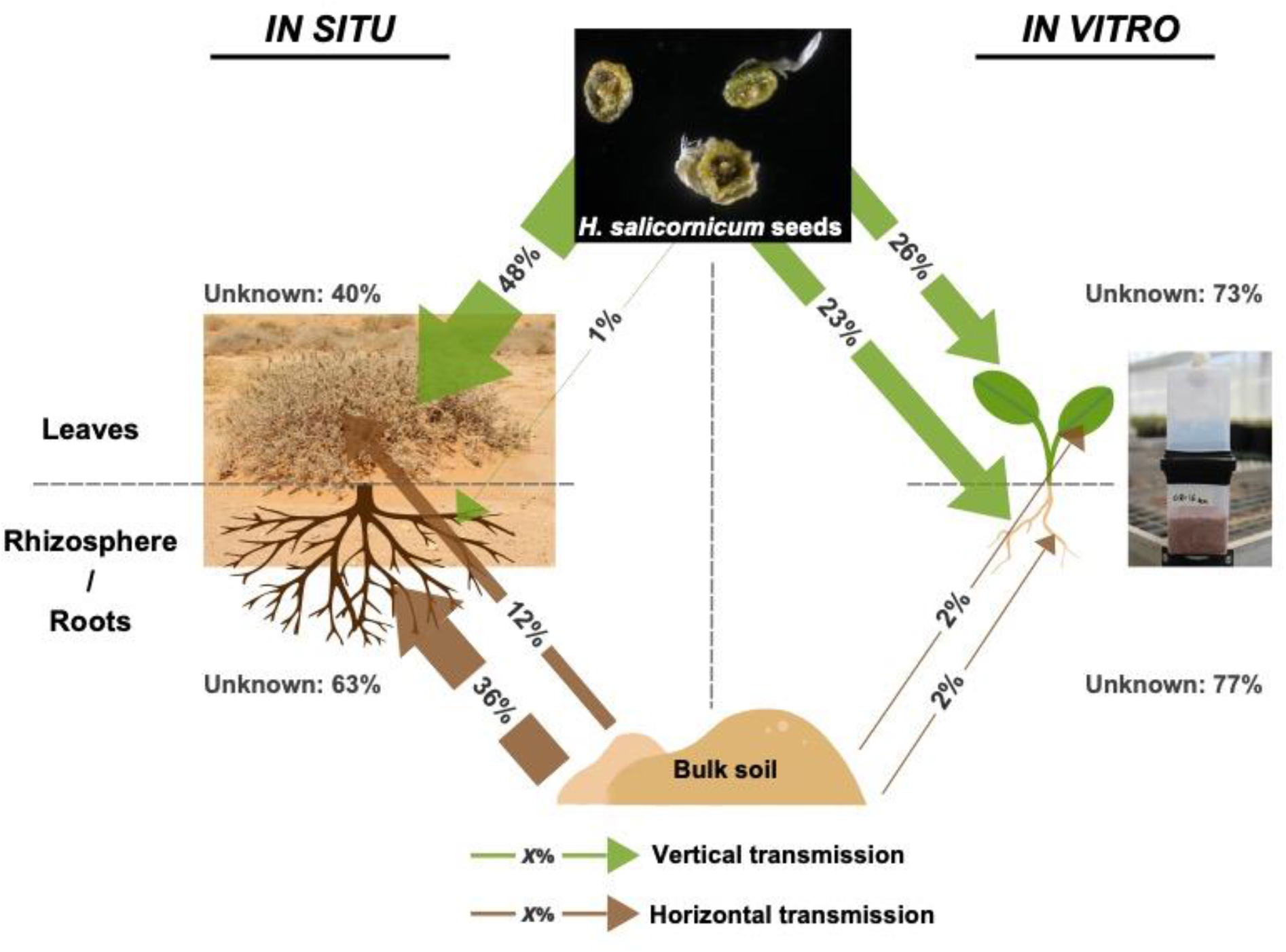
Root and rhizosphere mycobiota of adults *in situ* are mainly explained by the bulk soil (horizontal transmission), whereas leaf mycobiota observed *in situ* and leaf and root mycobiota of seedlings *in vitro* are mainly explained by seeds (vertical transmission).

#### Soil contributes to root and rhizosphere mycobiota *in situ*, while seeds contribute to leaf mycobiota *in situ* and leaf and root mycobiota *in vitro*

The source tracking algorithm *FEAST* and the identification of potentially transmitted OTUs revealed variable contributions of seeds (vertical transmission) and bulk soil (horizontal transmission) as potential fungal sources to *H. salicornicum* mycobiota according to the different compartments (Fig. 4). *In vitro*, both leaf and root mycobiota in the two substrate conditions (non- and autoclaved) were mainly explained by seed mycobiota (ranging from 16% to 33%; Fig.4a; Supp. Table 4), whereas the bulk soil contribution was very low (ranging from 0.1% for leaves in the non-autoclaved condition to 4.0% for leaves in the autoclaved condition). The contribution of seeds was significantly greater than that of bulk soil *in vitro* in both compartments and substrate conditions (with the exception of leaves of the non-autoclaved condition; Fig. 4a), suggesting a predominance of vertical transmission in seedlings *in vitro*. Both compartment and substrate condition variables had no significant effect on the proportions of the mycobiota explained by the source contributions *in vitro* (*p*>0.5 for both variables, *ANOVA*; Supp. Table 5; Supp. Fig. 6). In contrast, for adults *in situ*, we observed contrasted transmission pathways: the largest contribution of their roots and rhizosphere mycobiota is explained by the bulk soil (mean contribution of respectively 39 and 32%), which is significantly higher than the contribution explained by the seeds (respectively 2.0 and 0.8%; Fig. 4a; Supp. Table 4). Conversely, *in situ,* leaf mycobiota were mostly explained by the seeds (mean contribution of 48%) with a minor contribution from the bulk soil (12%).

Using our strict definition of potentially transmitted OTUs, we confirmed the prevalence of OTU sharing between seeds and seedlings: between 22 and 59 OTUs were shared between seeds and seedlings *in vitro*, representing 30 to 43% of their mycobiota (Fig. 4b, Supp. Table 4). In particular, *Mycosphaerella* and *Leucosporidium* OTUs were shared between seeds and roots/leaves in the two substrate conditions (Fig. 4b). Conversely, seedlings *in vitro* shared very few OTUs with the bulk soil (Fig. 4b), representing a null (or close to) contribution of their mycobiota (Supp. Table 4). This markedly contrasts with the rhizosphere and roots of adults *in situ* that shared a high number of OTUs with the bulk soil (respectively, 226 and 159; Fig. 4b), representing a large share of their mycobiota (43 and 40%, respectively; Supp. Table 4). Among candidates for transmission between bulk soil and rhizosphere/roots, we mainly identified OTUs belonging to the *Penicillium*, *Chaetomium*, and *Eutypella* genera (Fig. 4b). A very small number of OTUs were shared between seeds and rhizosphere/roots *in situ*, representing a small share of their mycobiota (Supp. Table 4). Bulk soil and leaves of adults *in situ* shared 46 OTUs, which represent 6.8% of the leaf mycobiota. This percentage is similar to the sharing between seeds and leaves *in situ* (54 OTUs, 7.9% of the mycobiota); this contrasts with the results from the source tracking analysis which indicated that the seeds contributed 48% of the leaf mycobiota *in situ* (Fig. 4a; Supp. Table 4). This apparent contradiction between the two methods was overcome when using our loose definition of potentially transmitted OTUs as we thus observed that some ubiquitous fungi (such as fungi from the genus *Alternaria* (Supp. Fig. 7) may be transmitted *in situ* by both seeds and bulk soil to rhizosphere, roots, and leaves. With this loose definition, potentially transmitted OTUs from seeds and bulk soil represented almost 40% of leaves *in situ* (Supp. Fig. 7).

Altogether, the source tracking algorithm and the identification of potentially transmitted OTUs were consistent, as the number of potentially transmitted OTUs (and their share of the mycobiota) was correlated (Supp. Fig. 8). They suggested a predominance of vertical transmission from seeds to seedlings *in vitro* (regardless of the compartment) and a predominance of horizontal transmission in soil compartments of adults *in situ*. Leaves of adults *in situ* displayed a mixed pattern, with a predominance of vertical transmission but a significant share of horizontal transmission.

## Discussion

We quantified the respective contributions of vertical (seeds) and horizontal (bulk soil) transmission pathways to the *H. salicornicum* mycobiota in seedlings *in vitro* and fully developed individuals *in situ*, for aerial and underground compartments. Roots and rhizosphere mycobiota of adult individuals *in situ* are mainly explained by bulk soil (horizontal transmission) with almost no contribution of seeds, whereas roots and leaves of seedlings *in vitro* are mainly explained by the seeds (vertical transmission) with almost no contribution from the soil (Fig. 5). Only leaves of adults *in situ* display a mixed pattern with a predominance of vertical transmission.

Summary of the estimated contribution of seeds and bulk soil contribution to *Haloxylon salicornicum* mycobiota in each compartment using the *FEAST* source tracking algorithm. Arrow width is proportional to the estimated contribution. For readability purposes, the contributions of bulk soil to roots and rhizosphere mycobiota of adults *in situ* were averaged as they are not significantly different (Fig. 4a). Similarly, as the contribution of both seeds and bulk soil to both leaf and root mycobiota in seedlings *in vitro* are not significantly different (Fig. 4a), they were averaged. The unexplained contribution to the mycobiota of adults *in situ* and seedlings *in vitro* is also reported.

### Vertical transmission is predominant in seedling mycobiota *in vitro* while bulk soil has a quasi-null influence

Both leaves and roots of seedlings germinated *in vitro* in (non-)autoclaved soil collected *in situ* display similar mycobiota composition resembling that of seeds, suggesting a predominance of vertical transmission. Due to a low germination rate and events of damping off in previous trials, we collected seedlings *in vitro* at an early development stage (2-leaf, 7 days). Despite a limited number of seedlings, we observed strong patterns across samples. First, the mycobiota of seedling roots and leaves *in vitro* were almost undifferentiated, with similar richness and community composition. Yet, compartment is known to be a strong factor of differentiation of the mycobiota, especially between above- and belowground compartments (Wearn *et al*., 2012; Martins *et al*., 2016; Harrison and Griffin, 2020), even though differences at juvenile stages may be lower as distance between compartments is reduced. Second, both substrate conditions (non- and autoclaved) used in the *in vitro* experiment resulted in similar mycobiota composition after 7 days, suggesting a low influence of this reservoir on seedling mycobiota. As evidenced by bipartite network analysis, seeds and seedlings (leaves and roots) tend to share a lot of OTUs, while bulk soil samples do not. This is supported by the source tracking analysis which estimated a quasi-null contribution of the soil to seedling mycobiota and by the low percentage of potentially transmitted OTUs from soil in seedling mycobiota (leaves and roots), in opposition to our initial hypothesis. On the contrary, seeds explain the largest part of both seedling root and leaf mycobiota (16 to 33% of the mycobiota is explained by seeds *in vitro*). Vertical transmission of the mycobiota (or part of it) may be favored in desert shrubs as their patchy distribution (fertility islands hypothesis) limits the dissemination of fungi to neighboring conspecific individuals. Our results are at odds with those of Rochefort *et al*. (2021): they observed that differences in initial soil conditions (such as differences in the initial quantity of soil microorganisms in the germination substrate) led to distinct fungal communities at 7 and 14 days in *Brassica napus* (Brassicaceae), suggesting that soil microbiota has a significantly greater influence on seedling mycobiota than seeds. These differences may be linked to differences in *in vitro* experimental settings: Rochefort et al. (2021) used gamma-irradiated soil inoculated with ‘active’ soil suspensions, a process which may lead to changes in mycobiota richness and composition and could in particular favor fungi with strong colonization capacities (Mesny *et al*., 2021). Other studies have shown that bacterial microbiota is vertically transmitted to seedlings by seeds (Moroenyane *et al*., 2021 on soybean; Walsh *et al*., 2021 on wheat) using inoculated seeds germinated *in vitro*. In our experiment, we used as substrate soil collected *in situ* which was unprocessed as much as possible before setting up the *in vitro* experiment. Though sampling may have altered the physiological condition and colonization capacity of soil fungi by breaking hyphae and limiting access to water, fungi in desert ecosystems have developed adaptations to the harsh conditions of these environments (sandy soil, nutrition depletion, low water availability, heat…), such as resistant spores, melanized hyphae, or biofilm formation with other organisms such as bacteria (Sterflinger *et al*., 2012; Ameen *et al*., 2021). We therefore hypothesize that soil sampling only had a limited effect on the colonization capacity of soil fungi in our experiment. These experimental biases could be overcome by sampling seedlings *in situ* (when possible) in addition to *in vitro* experiments.

Fungi potentially transmitted from seeds to seedlings were similar in the two compartments (roots and leaves) and substrate conditions (non- and autoclaved), and were mainly *Mycosphaerella asteroma* (ca. 10% of the seedling mycobiota). *Mycosphaerella* spp. have been mainly reported as plant pathogens (Hunter *et al*., 2011), yet we did not find any visual traces of plant diseases on seedlings at sampling time. Indeed, some *Mycosphaerella* species occur as symptomless endophytes (Kaneko *et al*., 2003; González-Teuber, 2016) and shift from pathogenic to endophytic, a behavior which is common among fungi (Selosse *et al*., 2018). These endophytic fungi may therefore be either harmless or latent pathogens. Other fungi belong to the yeast genera *Leucosporidium*, the genus *Vishniacozyma*, and *Aureobasidium* (*A*. *pullulans*), which has anti-fungal activities (Wachowska and Głowacka, 2014). Their roles as seed and seedling endophytes are poorly understood and deserve further attention.

A large share of seedling mycobiota *in vitro* is not explained by seeds or soil. Rarefaction curves of both sources (bulk soil and seed samples) tend to reach a plateau for most of the samples, suggesting that our sampling properly describes their fungal diversity. Furthermore, we used extraction and PCR negative controls along with stringent filtering in order to limit the presence of contaminants from molecular analysis. Unexplained fungal transmission may occur from other sources such as air (Zhou *et al*., 2021) and water - even though our experimental setup should limit these (Material & Methods – *In vitro* germination experiment).

### Rhizosphere and root mycobiota of adults *in situ* are mainly obtained by horizontal transmission, while leaf mycobiota are acquired by both vertical and horizontal transmission

Contrary to seedlings *in vitro*, adult individuals *in situ* display contrasted mycobiota composition according to the compartment studied. The main differences were observed between above- and belowground plant compartments (rhizosphere and roots vs. leaves and seeds). These observations are consistent with the source tracking analysis as both rhizosphere and root mycobiota are mainly explained by the bulk soil (39 and 33%, respectively), while the contribution of seeds is quasi-null. These results suggest that the mycobiota of belowground tissues of adults *in situ* are mainly transmitted horizontally, as hypothesized in this study. Potentially transmitted OTUs have similar taxonomy and are mainly affiliated to *Penicillium oxalicum* (ca. 10% of the root and rhizosphere mycobiota), *Eutypella* sp. (4%), Pleosporaceae sp. (3.5%), and *Chaetomium* sp. (3%). *P. oxalicum* is commonly found in the rhizosphere and was reported as a plant growth-promoting fungus which could limit the development of some pathogens such as *Fusarium* spp. (Murali and Amruthesh, 2015), while the others may be plant pathogens (esp. *Eutypella* sp.) or endophytes (such as *Chaetomium* sp.; FUNGuild online database, Nguyen *et al*., 2016).

Leaf mycobiota of adult individuals *in situ* display a mixed pattern of colonization. Their composition is similar to seeds which are the main contributor to their mycobiota (48%). This contribution is mainly supported by ubiquitous fungi such as *Alternaria consortialis* which represent a large share of leaf mycobiota *in situ* (10%) when using the loose definition of potentially transmitted OTUs. Notably, bulk soil has a non-null contribution to leaf mycobiota (12%), also characterized by the contribution of *A. consortialis*. Species from the *Alternaria* genus are common seed endophytes (Simonin *et al*., 2022) and have also been reported as widespread saprotroph and plant pathogens in leaves and shoots (Dang *et al*., 2015). *Alternaria* spp. have already been isolated from desert soils in the Arabian desert (Ameen *et al*., 2021) and our results suggest that they may have been recruited from both bulk soil and seeds *in situ*. Contribution from soil to phyllosphere mycobiota has already been shown for bacteria (Xiong *et al*., 2021), but the mechanisms of transmission from soil to phyllosphere are poorly described. Some authors suggest that soilborne microbes may be transmitted by air (Zhou *et al*., 2021) or within plant tissues (Vandenkoornhuyse *et al*., 2015) and indeed desert soils are poorly covered by vegetation, allowing transfer of dust to air. Our results support transmission of soil fungi to the phyllosphere, but do not allow us to draw conclusions regarding the pathways followed by these fungi. As for seedlings *in vitro*, a large share of *in situ* adult mycobiota is not explained by seeds or bulk soil. Contrary to the *in vitro* experiment, sources of contaminations *in situ* may be more diverse, including fungi dispersed by wind and small mammals (Zhou *et al*., 2021; Borgmann-Winter *et al*.), or even plant debris from the litter (Christian *et al*., 2017). Though fungi may be transmitted by wind (e.g., spores), the low density of plant individuals (in particular *H. salicornicum* individuals; fertility islands hypothesis) may limit their dispersion and therefore favor the withholding of vertically-transmitted fungi in *H. salicornicum* leaves, while the mycobiota of belowground tissues are more likely to be influenced by the environment (horizontal transmission).

### Vertical and horizontal transmission during microbiota assembly

Here, transmission from seeds may be pseudo-vertical as we investigated the total mycobiota (epiphyte and endophyte) of seeds collected on *H. salicornicum* individuals (before falling on the ground) and germinated them without surface sterilization. Vertical transmission *sensu stricto* describes the transmission of microorganisms strictly to the progeny, i.e., with no contamination from the environment (Bright and Bulgheresi, 2010; Truyens *et al*., 2015). However, seeds are exposed to several environmental contaminations over their life cycle from the early development stage to maturation and germination (Abdelfattah, *et al*., 2022; Nelson, 2018), resulting in modified microbiota/mycobiota which are also transmitted to seedlings. Once pseudo-vertical transmission is ensured, it may allow a partner fidelity sufficient to ensure the evolution of important functions for the host plant (Séne *et al*., 2018).

The contribution of seeds to *H. salicornicum* seedling root mycobiota *in vitro* was high, while bulk soil had a quasi-null contribution, an opposite pattern compared to that of adult root and rhizosphere mycobiota *in situ*. These opposite patterns may be linked not only to differences in life conditions (*in situ* vs. *in vitro*) but also to age, and reflect the juvenile mycobiota, which may also change during *H. salicornicum* development from the establishment of seedlings to adult individuals, as often reported (Houlden *et al*., 2008; Han *et al*., 2017; Gao *et al*., 2019; Liu and Howell, 2021). In the latter case, our results suggest a secondary colonization of roots and rhizosphere by soil fungi, replacing fungi vertically transmitted to seedlings. Plants can indeed recruit fungal partners actively by emitting signal molecules (Daguerre et al., 2020), while some fungi colonize compartments without any active recruitment from the plant (Gao et al., 2020), leading to changes in mycobiota composition with age (ecological recruitment process). As shown in previous studies, the initial microbiota may also impact the recruitment of fungi after germination by either limiting the development of other species or facilitating their establishment in plant tissues, a phenomenon called ‘priority effect’ (Ridout et al., 2019; Debray et al., 2022). Fungi pseudo-vertically transmitted may therefore play a crucial role in the mycobiota assembly by facilitating or limiting the colonization of other fungi from soil. Differences in horizontal and vertical transmission between *in situ* adults and *in vitro* seedlings are less contrasted in leaf mycobiota as in both cases, seeds (vertical transmission) are the main contributor to their mycobiota. However, we identified a significant share of horizontal transmission from the soil to leaves of adults *in situ* but none in seedlings *in vitro*. Furthermore, fungi potentially transmitted from seeds to leaves differ between adults *in situ* and seedlings *in vitro*. Again, these differences may be linked to differences between environmental conditions, but also to changes in mycobiota composition of aerial compartments during plant development (Maignien *et al*., 2014). Understanding whether fungi are transmitted horizontally or vertically has crucial ecological implications over short timescales, but may also help us to understand phenomena occurring over long timescales such as phylosymbiosis patterns (i.e., phylogenetically close individuals tend to associate preferentially with related hosts), which have been observed in plants with fungi (Perez-Lamarque *et al*., 2020) and bacteria (Abdelfattah, *et al*., 2022), interactions suggesting preferential associations between plants and fungi and/or bacteria. Some authors suggest that such evolutionary patterns are testimony of microbiota transmission from one generation to the other (Abdelfattah, et al., 2022), reflecting the relevance of our findings, although host filtering from the environment may be the main mechanism at stake (Mazel et al., 2018).

## Conclusion

Through a combination of *in situ* sampling and *in vitro* experiment, we have described and quantified vertical (from seeds) and horizontal transmission (from bulk soil) of *H. salicornicum* mycobiota (Fig. 5). We show *in vitro* that the mycobiota are partially transmitted to seedlings by seeds (vertical transmission) while soil does not influence their mycobiota, contrary to our hypothesis and previous results. The mycobiota of belowground compartments of *in situ* adult individuals display an opposite pattern as the contribution of seeds is almost null: rhizosphere and root mycobiota are mainly explained by the bulk soil (horizontal transmission). Leaves of adult individuals display a mixed pattern as their mycobiota are mainly explained by seeds, with a significant contribution of bulk soil. However, fungi transmitted from seeds to leaves differ between adults and seedlings. Differences in transmission pathways between adults *in situ* and seedlings *in vitro* may be linked to differences in experimental designs, but also to differences between developmental stages. Taking dynamic changes in the mycobiota and the transmission pathways into account would improve our understanding of mycobiota assembly. Finally, we show that some fungi are specifically shared between a source and a compartment (e.g., bulk soil and roots sharing *P. oxalicum*), while some ubiquitous fungi (such as *A*. *consortialis*) may be transmitted by both seeds and soil. The role of these fungi needs further attention, in particular, to decipher whether fungi transmitted by seeds are beneficial or deleterious for the host.

## Supporting information

Supplemental Informations

## Acknowledgments

We would like to thank Dr. Sami YOUSSEF and the company Valorhiz for providing the *Haloxylon salicornicum* seeds.

## References

Abdelfattah, A., Tack, A.J.M., Lobato, C., Wassermann, B., and Berg, G. (2022) From seed to seed: the role of microbial inheritance in the assembly of the plant microbiome. Trends Microbiol.

Abdelfattah, A., Tack, A.J.M., Wasserman, B., Liu, J., Berg, G., Norelli, J., et al. (2022) Evidence for host–microbiome co-evolution in apple. New Phytol 234: 2088–2100.

Abdelfattah, A., Wisniewski, M., Schena, L., and Tack, A.J.M. (2021) Experimental evidence of microbial inheritance in plants and transmission routes from seed to phyllosphere and root. Environ Microbiol 23: 2199–2214.

Al Salameen, F., Habibi, N., Kumar, V., Al Amad, S., Dashti, J., Talebi, L., and Al Doaij, B. (2018) Genetic diversity and population structure of Haloxylon salicornicum moq. in Kuwait by ISSR markers. PLOS ONE 13: e0207369.

Ameen, F., Stephenson, S.L., Al Nadhari, S., and Yassin, M.A. (2021) A review of fungi associated with Arabian desert soils. Nova Hedwig 173–195.

Barret, M., Briand, M., Bonneau, S., Préveaux, A., Valière, S., Bouchez, O., et al. (2015) Emergence Shapes the Structure of the Seed Microbiota. Appl Environ Microbiol 81: 1257–1266.

Berg, G., Rybakova, D., Fischer, D., Cernava, T., Vergès, M.-C.C., Charles, T., et al. (2020) Microbiome definition re-visited: old concepts and new challenges. Microbiome 8: 103.

Blüthgen, Nico, Menzel, F., and Blüthgen, Nils (2006) Measuring specialization in species interaction networks. BMC Ecol 6: 9.

Bonito, G., Reynolds, H., Robeson II, M.S., Nelson, J., Hodkinson, B.P., Tuskan, G., et al. (2014) Plant host and soil origin influence fungal and bacterial assemblages in the roots of woody plants. Mol Ecol 23: 3356–3370.

Borgmann-Winter, B.W., Stephens, R.B., Anthony, M.A., Frey, S.D., D’Amato, A.W., and Rowe, R.J. Wind and small mammals are complementary fungal dispersers. Ecology n/a: e4039.

Brigham, L.M., Bueno de Mesquita, C.P., Spasojevic, M.J., Farrer, E.C., Porazinska, D.L., Smith, J.G., et al. (2023) Drivers of bacterial and fungal root endophyte communities: understanding the relative influence of host plant, environment, and space. FEMS Microbiol Ecol 99: fiad034.

Bright, M. and Bulgheresi, S. (2010) A complex journey: transmission of microbial symbionts. Nat Rev Microbiol 8: 218–230.

Bulgarelli, D., Schlaeppi, K., Spaepen, S., van Themaat, E.V.L., and Schulze-Lefert, P. (2013) Structure and Functions of the Bacterial Microbiota of Plants. Annu Rev Plant Biol 64: 807–838.

Cai, L., Kreft, H., Taylor, A., Denelle, P., Schrader, J., Essl, F., et al. (2023) Global models and predictions of plant diversity based on advanced machine learning techniques. New Phytol 237: 1432–1445.

Christian, N., Herre, E.A., Mejia, L.C., and Clay, K. (2017) Exposure to the leaf litter microbiome of healthy adults protects seedlings from pathogen damage. Proc R Soc B Biol Sci 284: 20170641.

Csardi, G. and Nepusz, T. (2006) The igraph software package for complex network research. InterJournal Complex Syst 1695: 1–9.

Daguerre, Y., Basso, V., Hartmann-Wittulski, S., Schellenberger, R., Meyer, L., Bailly, J., et al. (2020) The mutualism effector MiSSP7 of Laccaria bicolor alters the interactions between the poplar JAZ6 protein and its associated proteins. Sci Rep 10: 20362.

Dang, H.X., Pryor, B., Peever, T., and Lawrence, C.B. (2015) The Alternaria genomes database: a comprehensive resource for a fungal genus comprised of saprophytes, plant pathogens, and allergenic species. BMC Genomics 16: 239.

Debray, R., Herbert, R.A., Jaffe, A.L., Crits-Christoph, A., Power, M.E., and Koskella, B. (2022) Priority effects in microbiome assembly. Nat Rev Microbiol 20: 109–121.

Douglas, A.E. and Werren, J.H. (2016) Holes in the Hologenome: Why Host-Microbe Symbioses Are Not Holobionts. mBio 7: e02099–15.

Fruchterman, T.M.J. and Reingold, E.M. (1991) Graph drawing by force-directed placement. Softw Pract Exp 21: 1129–1164.

Gao, C., Montoya, L., Xu, L., Madera, M., Hollingsworth, J., Purdom, E., et al. (2020) Fungal community assembly in drought-stressed sorghum shows stochasticity, selection, and universal ecological dynamics. Nat Commun 11: 34.

Gao, C., Montoya, L., Xu, L., Madera, M., Hollingsworth, J., Purdom, E., et al. (2019) Strong succession in arbuscular mycorrhizal fungal communities. ISME J 13: 214–226.

Gloor, G.B., Macklaim, J.M., Pawlowsky-Glahn, V., and Egozcue, J.J. (2017) Microbiome Datasets Are Compositional: And This Is Not Optional. Front Microbiol 8:.

González-Teuber, M. (2016) The defensive role of foliar endophytic fungi for a South American tree. AoB PLANTS 8: plw050.

de Graaff, M.-A., Throop, H.L., Verburg, P.S.J., Arnone, J.A., and Campos, X. (2014) A Synthesis of Climate and Vegetation Cover Effects on Biogeochemical Cycling in Shrub-Dominated Drylands. Ecosystems 17: 931–945.

Gundel, P.E., Rudgers, J.A., and Ghersa, C.M. (2011) Incorporating the process of vertical transmission into understanding of host–symbiont dynamics. Oikos 120: 1121–1128.

Han, L.-L., Wang, J.-T., Yang, S.-H., Chen, W.-F., Zhang, L.-M., and He, J.-Z. (2017) Temporal dynamics of fungal communities in soybean rhizosphere. J Soils Sediments 17: 491–498.

Hardoim, P.R., Hardoim, C.C.P., van Overbeek, L.S., and van Elsas, J.D. (2012) Dynamics of Seed-Borne Rice Endophytes on Early Plant Growth Stages. PLOS ONE 7: e30438.

Harrison, J.G. and Griffin, E.A. (2020) The diversity and distribution of endophytes across biomes, plant phylogeny and host tissues: how far have we come and where do we go from here? Environ Microbiol 22: 2107–2123.

van der Heijden, M.G.A., Martin, F.M., Selosse, M.-A., and Sanders, I.R. (2015) Mycorrhizal ecology and evolution: the past, the present, and the future. New Phytol 205: 1406–1423.

Hodgson, S., de Cates, C., Hodgson, J., Morley, N.J., Sutton, B.C., and Gange, A.C. (2014) Vertical transmission of fungal endophytes is widespread in forbs. Ecol Evol 4: 1199–1208.

Hosseyni Moghaddam, M.S., Safaie, N., Soltani, J., and Hagh-Doust, N. (2021) Desert-adapted fungal endophytes induce salinity and drought stress resistance in model crops. Plant Physiol Biochem 160: 225–238.

Houlden, A., Timms-Wilson, T.M., Day, M.J., and Bailey, M.J. (2008) Influence of plant developmental stage on microbial community structure and activity in the rhizosphere of three field crops. sFEMS Microbiol Ecol 65: 193–201.

Hunter, G.C., Crous, P.W., Carnegie, A.J., Burgess, T.I., and Wingfield, M.J. (2011) Mycosphaerella and Teratosphaeria diseases of Eucalyptus; easily confused and with serious consequences. Fungal Divers 50: 145–166.

Intergovernmental Panel on Climate Change ed. (2022) Desertification. In Climate Change and Land: IPCC Special Report on Climate Change, Desertification, Land Degradation, Sustainable Land Management, Food Security, and Greenhouse Gas Fluxes in Terrestrial Ecosystems. Cambridge: Cambridge University Press, pp. 249–344.

Johnston-Monje, D., Lundberg, D.S., Lazarovits, G., Reis, V.M., and Raizada, M.N. (2016) Bacterial populations in juvenile maize rhizospheres originate from both seed and soil. Plant Soil 405: 337–355.

Kaneko, R., Kakishima, M., and Tokumasu, S. (2003) The seasonal occurrence of endophytic fungus, Mycosphaerella buna, in Japanese beech, Fagus crenata. Mycoscience 44: 277–281.

Klaedtke, S., Jacques, M.-A., Raggi, L., Préveaux, A., Bonneau, S., Negri, V., et al. (2016) Terroir is a key driver of seed-associated microbial assemblages. Environ Microbiol 18: 1792–1804.

Legendre, P. and Gallagher, E.D. (2001) Ecologically meaningful transformations for ordination of species data. Oecologia 129: 271–280.

Li, X., He, X., Hou, L., Ren, Y., Wang, S., and Su, F. (2018) Dark septate endophytes isolated from a xerophyte plant promote the growth of Ammopiptanthus mongolicus under drought condition. Sci Rep 8: 7896.

Liu, D. and Howell, K. (2021) Community succession of the grapevine fungal microbiome in the annual growth cycle. Environ Microbiol 23: 1842–1857.

Lundberg, D.S., Lebeis, S.L., Paredes, S.H., Yourstone, S., Gehring, J., Malfatti, S., et al. (2012) Defining the core Arabidopsis thaliana root microbiome. Nature 488: 86–90.

Maignien, L., DeForce, E.A., Chafee, M.E., Eren, A.M., and Simmons, S.L. (2014) Ecological Succession and Stochastic Variation in the Assembly of Arabidopsis thaliana Phyllosphere Communities. mBio 5: e00682–13.

Martin, M. (2011) Cutadapt removes adapter sequences from high-throughput sequencing reads. EMBnet.journal 17: 10.

Martins, F., Pereira, J.A., Bota, P., Bento, A., and Baptista, P. (2016) Fungal endophyte communities in above- and belowground olive tree organs and the effect of season and geographic location on their structures. Fungal Ecol 20: 193–201.

Maurice, K., Laurent-Webb, L., Dehail, A., Bourceret, A., Boivin, S., Boukcim, H., et al. (2023) Fertility Islands, Keys to the Establishment of Plant and Microbial Diversity in a Highly Alkaline Hot Desert. DOI.x:10.2139/ssrn.4341673.

Mazel, F., Davis, K.M., Loudon, A., Kwong, W.K., Groussin, M., and Parfrey, L.W. (2018) Is Host Filtering the Main Driver of Phylosymbiosis across the Tree of Life? mSystems 3: e00097–18.

McKnight, D.T., Huerlimann, R., Bower, D.S., Schwarzkopf, L., Alford, R.A., and Zenger, K.R. (2019) Methods for normalizing microbiome data: An ecological perspective. Methods Ecol Evol 10: 389–400.

McMurdie, P.J. and Holmes, S. (2013) phyloseq: An R Package for Reproducible Interactive Analysis and Graphics of Microbiome Census Data. PLOS ONE 8: e61217.

Mesny, F., Miyauchi, S., Thiergart, T., Pickel, B., Atanasova, L., Karlsson, M., et al. (2021) Genetic determinants of endophytism in the Arabidopsis root mycobiome. Nat Commun 12: 7227.

Moroenyane, I., Tremblay, J., and Yergeau, É. (2021) Soybean Microbiome Recovery After Disruption is Modulated by the Seed and Not the Soil Microbiome. Phytobiomes J 5: 418–431.

Murali, M. and Amruthesh, K.N. (2015) Plant Growth-promoting Fungus Penicillium oxalicum Enhances Plant Growth and Induces Resistance in Pearl Millet Against Downy Mildew Disease. J Phytopathol 163: 743–754.

Nelson, E.B. (2018) The seed microbiome: Origins, interactions, and impacts. Plant Soil 422: 7–34.

Nguyen, N.H., Song, Z., Bates, S.T., Branco, S., Tedersoo, L., Menke, J., et al. (2016) FUNGuild: An open annotation tool for parsing fungal community datasets by ecological guild. Fungal Ecol 20: 241–248.

Nilsson, R.H., Larsson, K.-H., Taylor, A.F.S., Bengtsson-Palme, J., Jeppesen, T.S., Schigel, D., et al. (2019) The UNITE database for molecular identification of fungi: handling dark taxa and parallel taxonomic classifications. Nucleic Acids Res 47: D259–D264.

Ochoa-Hueso, R., Eldridge, D.J., Delgado-Baquerizo, M., Soliveres, S., Bowker, M.A., Gross, N., et al. (2018) Soil fungal abundance and plant functional traits drive fertile island formation in global drylands. J Ecol 106: 242–253.

Oksanen, J., Blanchet, F.G., Kindt, R., Legendre, P., Minchin, P.R., O’hara, R.B., et al. (2013) Package ‘vegan.’ Community Ecol Package Version 2: 1–295.

Op De Beeck, M., Lievens, B., Busschaert, P., Declerck, S., Vangronsveld, J., and Colpaert, J.V. (2014) Comparison and Validation of Some ITS Primer Pairs Useful for Fungal Metabarcoding Studies. PLoS ONE 9: e97629.

Perez-Lamarque, B., Krehenwinkel, H., Gillespie, R.G., and Morlon, H. (2022) Limited Evidence for Microbial Transmission in the Phylosymbiosis between Hawaiian Spiders and Their Microbiota. mSystems 7: e01104-21.

Perez-Lamarque, B., Laurent-Webb, L., Bourceret, A., Maillet, L., Bik, F., Cartier, D., et al. (2023) Fungal microbiomes associated with Lycopodiaceae during ecological succession. Environ Microbiol Rep 15: 109–118.

Perez-Lamarque, B., Petrolli, R., Strullu-Derrien, C., Strasberg, D., Morlon, H., Selosse, M.-A., and Martos, F. (2022) Structure and specialization of mycorrhizal networks in phylogenetically diverse tropical communities. Environ Microbiome 17: 38.

Perez-Lamarque, B., Selosse, M.-A., Öpik, M., Morlon, H., and Martos, F. (2020) Cheating in arbuscular mycorrhizal mutualism: a network and phylogenetic analysis of mycoheterotrophy. New Phytol 226: 1822–1835.

Petrolli, R., Augusto Vieira, C., Jakalski, M., Bocayuva, M.F., Vallé, C., Cruz, E.D.S., et al. (2021) A fine-scale spatial analysis of fungal communities on tropical tree bark unveils the epiphytic rhizosphere in orchids. New Phytol 231: 2002–2014.

Prăvălie, R. (2016) Drylands extent and environmental issues. A global approach. Earth-Sci Rev 161: 259–278.

R Core Team (2023) R: A Language and Environment for Statistical Computing, Vienna, Austria: R Foundation for Statistical Computing.

Rathore, V.S., Singh, J.P., Bhardwaj, S., Nathawat, N.S., Kumar, M., and Roy, M.M. (2015) Potential of Native Shrubs Haloxylon salicornicum and Calligonum Polygonoides for Restoration of Degraded Lands in Arid Western Rajasthan, India. Environ Manage 55: 205–216.

Ridout, M.E., Schroeder, K.L., Hunter, S.S., Styer, J., and Newcombe, G. (2019) Priority effects of wheat seed endophytes on a rhizosphere symbiosis. Symbiosis 78: 19–31.

Robinson, R.J., Fraaije, B.A., Clark, I.M., Jackson, R.W., Hirsch, P.R., and Mauchline, T.H. (2016) Wheat seed embryo excision enables the creation of axenic seedlings and Koch’s postulates testing of putative bacterial endophytes. Sci Rep 6: 25581.

Rochefort, A., Simonin, M., Marais, C., Guillerm-Erckelboudt, A.-Y., Barret, M., and Sarniguet, A. (2021) Transmission of Seed and Soil Microbiota to Seedling. *mSystems* 6: e00446-21.

Rodríguez, C.E., Antonielli, L., Mitter, B., Trognitz, F., and Sessitsch, A. (2020) Heritability and Functional Importance of the Setaria viridis Bacterial Seed Microbiome. Phytobiomes J 4: 40– 52.

Rodriguez Estrada, A.E., Jonkers, W., Corby Kistler, H., and May, G. (2012) Interactions between Fusarium verticillioides, Ustilago maydis, and Zea mays: An endophyte, a pathogen, and their shared plant host. Fungal Genet Biol 49: 578–587.

Rodriguez, R.J., Jr, J.F.W., Arnold, A.E., and Redman, R.S. (2009) Fungal endophytes: diversity and functional roles. New Phytol 182: 314–330.

Rognes, T., Flouri, T., Nichols, B., Quince, C., and Mahé, F. (2016) VSEARCH: a versatile open source tool for metagenomics. PeerJ 4: e2584.

Schlesinger, W.H. and Pilmanis, A.M. (1998) Plant-soil Interactions in Deserts. Biogeochemistry 42: 169–187.

Schneider-Maunoury, L., Deveau, A., Moreno, M., Todesco, F., Belmondo, S., Murat, C., et al. (2020) Two ectomycorrhizal truffles, Tuber melanosporum and T. aestivum, endophytically colonise roots of non-ectomycorrhizal plants in natural environments. New Phytol 225: 2542–2556.

Selosse, M.-A., Baudoin, E., and Vandenkoornhuyse, P. (2004) Symbiotic microorganisms, a key for ecological success and protection of plants. C R Biol 327: 639–648.

Selosse, M.-A., Bessis, A., and Pozo, M.J. (2014) Microbial priming of plant and animal immunity: symbionts as developmental signals. Trends Microbiol 22: 607–613.

Séne, S., Selosse, M.-A., Forget, M., Lambourdière, J., Cissé, K., Diédhiou, A.G., et al. (2018) A pantropically introduced tree is followed by specific ectomycorrhizal symbionts due to pseudo-vertical transmission. ISME J 12: 1806–1816.

Shade, A., Jacques, M.-A., and Barret, M. (2017) Ecological patterns of seed microbiome diversity, transmission, and assembly. Curr Opin Microbiol 37: 15–22.

Shenhav, L., Thompson, M., Joseph, T.A., Briscoe, L., Furman, O., Bogumil, D., et al. (2019) FEAST: fast expectation-maximization for microbial source tracking. Nat Methods 16: 627–632.

Simonin, M., Briand, M., Chesneau, G., Rochefort, A., Marais, C., Sarniguet, A., and Barret, M. (2022) Seed microbiota revealed by a large-scale meta-analysis including 50 plant species. New Phytol 234: 1448–1463.

Singh, J.P., Rathore, V.S., and Roy, M.M. (2015) Notes about Haloxylon salicornicum (Moq.) Bunge ex Boiss., a promising shrub for arid regions. Genet Resour Crop Evol 62: 451–463.

Smith, S.E. and Read, D.J. (2009) Mycorrhizal symbiosis, 3. ed., Repr. Amsterdam: Elsevier/Acad. Press.

Sterflinger, K., Tesei, D., and Zakharova, K. (2012) Fungi in hot and cold deserts with particular reference to microcolonial fungi. Fungal Ecol 5: 453–462.

Susana Rivera, C., Eugenia Venturini, M., Oria, R., and Blanco, D. (2011) Selection of a decontamination treatment for fresh Tuber aestivum and Tuber melanosporum truffles packaged in modified atmospheres. FOOD CONTROL 22: 626–632.

Tedersoo, L., Bahram, M., Põlme, S., Kõljalg, U., Yorou, N.S., Wijesundera, R., et al. (2014) Global diversity and geography of soil fungi. Science 346: 1256688.

Tedersoo, L., Bahram, M., Zinger, L., Nilsson, R.H., Kennedy, P.G., Yang, T., et al. (2022) Best practices in metabarcoding of fungi: From experimental design to results. Mol Ecol 31: 2769–2795.

Tintjer, T., Leuchtmann, A., and Clay, K. (2008) Variation in horizontal and vertical transmission of the endophyte Epichloë elymi infecting the grass Elymus hystrix. New Phytol 179: 236–246.

Trivedi, P., Leach, J.E., Tringe, S.G., Sa, T., and Singh, B.K. (2020) Plant–microbiome interactions: from community assembly to plant health. Nat Rev Microbiol 18: 607–621.

Truyens, S., Weyens, N., Cuypers, A., and Vangronsveld, J. (2015) Bacterial seed endophytes: genera, vertical transmission and interaction with plants. Environ Microbiol Rep 7: 40–50.

Vandenkoornhuyse, P., Quaiser, A., Duhamel, M., Van, A.L., and Dufresne, A. (2015) The importance of the microbiome of the plant holobiont. New Phytol 206: 1196–1206.

Vannier, N., Mony, C., Bittebiere, A.-K., Michon-Coudouel, S., Biget, M., and Vandenkoornhuyse, P. (2018) A microorganisms’ journey between plant generations. Microbiome 6: 79.

Wachowska, U. and Głowacka, K. (2014) Antagonistic interactions between Aureobasidium pullulans and Fusarium culmorum, a fungal pathogen of winter wheat. BioControl 59: 635–645.

Walsh, C.M., Becker-Uncapher, I., Carlson, M., and Fierer, N. (2021) Variable influences of soil and seed-associated bacterial communities on the assembly of seedling microbiomes. ISME J 15: 2748–2762.

Wearn, J.A., Sutton, B.C., Morley, N.J., and Gange, A.C. (2012) Species and organ specificity of fungal endophytes in herbaceous grassland plants. J Ecol 100: 1085–1092.

White, T.J., Bruns, T., Lee, S., and Taylor, J. (1990) AMPLIFICATION AND DIRECT SEQUENCING OF FUNGAL RIBOSOMAL RNA GENES FOR PHYLOGENETICS. In PCR Protocols. Elsevier, pp. 315– 322.

Wilkinson, D.M. (1997) The Role of Seed Dispersal in the Evolution of Mycorrhizae. Oikos 78: 394– 396.

Wilson, D. (1995) Endophyte: The Evolution of a Term, and Clarification of Its Use and Definition. Oikos 73: 274.

Wolf, D.C., Dao, T.H., Scott, H.D., and Lavy, T.L. (1989) Influence of Sterilization Methods on Selected Soil Microbiological, Physical, and Chemical Properties. J Environ Qual 18: 39–44.

Xiong, C., Zhu, Y.-G., Wang, J.-T., Singh, B., Han, L.-L., Shen, J.-P., et al. (2021) Host selection shapes crop microbiome assembly and network complexity. New Phytol 229: 1091–1104.

Yakti, W., Kovács, G.M., Vági, P., and Franken, P. (2018) Impact of dark septate endophytes on tomato growth and nutrient uptake. Plant Ecol Divers 11: 637–648.

Zhou, S.-Y.-D., Li, H., Giles, M., Neilson, R., Yang, X., and Su, J. (2021) Microbial Flow Within an Air-Phyllosphere-Soil Continuum. Front Microbiol 11:.

