## Supplemental Informations for "Seed or soil: tracing back the plant mycobiota primary sources"

### Supplementary Methods (1-3)

**<u>Supplementary Methods 1:</u> Library preparation**

We amplified the ITS2 region of the rDNA operon using ITS86F-ITS4 primers (White *et al.*, 1990; Op De Beeck *et al.*, 2014). PCR reactions were performed using the Thermo Scientific Phusion™ High-Fidelity DNA Polymerase (ThermoFisher Scientific, USA) in 20 µL of reaction mix containing: 10 µL of Buffer Master mix (containing the Taq Polymerase), 0.5 µL of DMSO, 4.5 µL of DNAse free water, 3 µL of DNA, 1 µL of Forward + 1 µL of Reverse primers (10 µM). PCR conditions were as follows: initial denaturation for 10 min, 35 cycles each composed of denaturation at 94°C for 10s, annealing at 55°C for 10s and extension at 72°C for 10s with a final extension at 72°C for 7 min.

PCR products were purified using Agencourt AMPure XP beads (Beckman Coulter Inc., Indianapolis, IN, USA). PCR products were mixed with AMPure beads solution in a 1:1 (v:v) ratio and placed on a magnetic rack to remove the supernatant. Magnetic beads were then rinsed with ethanol in excess before and DNA was eluted in 70 µL of EB buffer (Qiagen, Germany).

All purified PCR products concentrations were measured with PicoGreen (Quant-iT™ PicoGreen™ dsDNA Assay Kits, ThermoFisher Scientific, USA). PCR products from samples of bulk soil, rhizosphere and roots *in situ* were mixed in one equimolar pool. Samples of leaves, seeds, and seedlings *in vitro* were pooled in another equimolar pool. The pools were purified twice with AMPure XP beads and pooled together in equimolar amounts (Beckman Coulter Inc., Indianapolis, IN, USA). MetaFast library preparation and sequencing were performed by Fasteris SA (Switzerland) on an Illumina platform using the 2x250 pb Miseq technology.

**<u>Supplementary Methods 2:</u> Bio-informatic analysis**

A pipeline based on VSEARCH (Rognes *et al.*, 2016) and available in GitHub (https://github.com/BPerezLamarque/Scripts/) was used for data processing (Petrolli *et al.*, 2021; Perez-Lamarque *et al.*, 2022, 2023). Paired-end reads were merged (*--fastq_mergepairs* function, default parameters) and quality checked (*--fastq_maxns* 0, *--fastq_maxee* 2). Merged reads were then demultiplexed using *cutadapt* (Martin, 2011) with 0 error accepted in primers or tags sequences. Reads from all samples were dereplicated (*--derep_fulllength)*. ITS2 sequences were clustered as classical 97% sequence similarity Operational Taxonomic Units (OTUs) using VSEARCH (--*cluster*_size) as advised in Tedersoo et al. (2022). All sequences were checked for the presence of chimeras (--*uchime_denovo*). The taxonomy of fungi was assigned with VSEARCH against the UNITE v9.0 database (Nilsson *et al.*, 2019). Reads were filtered in order to keep only non-chimeric sequences of > 200 pb and with a total abundance of at least 10.

We used the *decontam* algorithm (Susana Rivera *et al.*, 2011) to remove potential contaminants in the datasets, using both algorithms (*prevalence* and *frequency*). First, we used the *prevalence* algorithm using the extraction and PCR controls corresponding to each sequencing run with a stringent threshold of 0.5. We then used the *frequency* algorithm using the default threshold. A few more filters were applied to all datasets: we removed samples less than 1 000 fungal reads, OTUs with less than 5 reads per sample and OTUs representing less than 0.5% of the reads per sample.

In order to compute UniFrac distances, we reconstructed the fungal phylogeny as in Perez-Lamarque *et al.* (2022): sequences were aligned with MAFFT (Katoh and Standley, 2013) and trimmed using trimAl (Capella-Gutiérrez *et al.*, 2009). We then constructed a maximum-likelihood tree using IQ-TREE (Nguyen *et al.*, 2015) with 1,000 SH-aLRT and ultrafast bootstraps.

After filtering, we obtained the mycobiota composition of 259 samples (Supp. Table 1), with a mean sequencing depth of 21 013 reads (ranging from 1 041 to 130 227; Supp. Fig. 2).

**Supplementary Methods 3: Statistical analysis**

OTUs tables were processed using the *phyloseq* package (McMurdie and Holmes, 2013) in R (R Core Team, 2023).

*Alpha-diversity***.** In order to test differences between experimental designs (*in situ* vs. *in vitro*), compartments (bulk soil, rhizosphere, roots, leaves and seeds) and substrate condition *in vitro* (non- vs. autoclaved) for richness (Chao1 index) and diversity (Shannon index) we used linear regressions and Tukey's post hoc pairwise test. For each index, we ran two models: (i) first, with all samples to test for differences between *in vitro* and *in situ* samples and between compartments using the following model: *alpha-diversity index ~ exp. design * compartment* (where *exp. design* corresponds to *in situ* or *in vitro*) and (ii) with only leaves and roots of individuals germinated *in vitro* to test differences between compartments and between the two substrate condition (non- and autoclaved bulk soil) using the following model: *alpha-diversity index ~ compartment * substrate condition*.

*Bipartite networks***.** We built weighted bipartite networks using the *igraph* R package (Csardi and Nepusz, 2006). Nodes represent OTUs and samples, and edges the relative abundance of an OTU in a sample. We build a total network (all samples) and a sub-network with *in vitro* samples, bulk soil used for the germination experiment and seeds. We also built one network by compartment *in situ* and one network by compartment x substrate condition combinations *in vitro*. For each network, we only kept OTUs representing more than 0.5% of the reads in each sample. In order to test for the significance of the connectance and $H_2$' network specialization (Blüthgen *et al.*, 2006), we constructed 100 randomized marginal networks (i.e., networks permuted from the original network keeping original marginal sums) for each network (*r2dtable*; Blüthgen et al., 2006; Dormann et al., 2008) and computed their connectance and $H_2$'. Connectance was considered significantly lower from the null models if less than 2.5% of the randomized networks had a superior or equal connectance. Similarly, specialization was considered significantly superior from null models if less than 2.5% of the randomized networks had a lower or equal specialization.

*Source tracking analysis***.** In order to estimate the respective contributions of seeds and bulk soil to the *H. salicornicum* microbiota, we used the *FEAST* source tracking algorithm developed by Shenhav *et al.* (2019) and implemented in R. This algorithm takes as inputs microbial communities to explain (the ‘sinks’) and potential source environments (the ‘sources’) to estimate the fraction of the sinks explained by the sources. The algorithm also reports an unexplained fraction referred to as the ‘unknown’ source. Here, seeds and bulk soil samples were defined as sources whereas roots, rhizosphere and leaves were defined as sinks. We ran the procedure twice: (i) first on samples *in situ* and a second time (ii) on a subset of samples with samples from the *in vitro* experiment, bulk soils samples used for the experiment (see *Sampling*) and seeds. Proportions were transformed using the semi-parametric *ordered quantile normalization* (ORQ; Peterson & Cavanaugh, 2020) to reach normality. We ran the following linear regressions: (i) ORQ(*proportion*) ~ *Source* * *compartment* for the *in-situ* dataset, where *Source* is either seeds, bulk soil or an unknown source and (ii) ORQ(*proportion*) ~ *Source* * *condition* * *compartment* for the sub-dataset. We used Tukey’s post-hoc multiple comparisons test for pairwise comparisons.

#### Supplementary Tables (1-6)

**Supplementary Table 1: Soil and *H. salicornicum* samples collected and successfully processed.**

For samples *in vitro*, the left number of the brackets is the number of samples in the autoclaved condition, and the right number (underlined) is the number of samples in the non-autoclaved condition.

[1] Data from Maurice *et al.* (2023).

[2] New data from this study.

| | | **Bulk soil** | **Rhizosphere** | **Roots** | **Leaves** | **Seeds** |
|---|---|---|---|---|---|---|
| **Collected samples** | ***In situ*** | 25 [1] | 65 [1] | 65 [1] | 65 [2] | 20 (pools of 9 seeds) [2] |
| | ***In vitro*** | | | 20 (13 + 7) [2] | 20 (13 + 7) [2] | |
| **Successfully processed samples** | ***In situ*** | 24 [1] | 63 [1] | 59 [1] | 61 [2] | 13 (pools of 9 seeds) [2] |
| | ***In vitro*** | | | 19 (13 + 6) [2] | 20 (13 + 7) [2] | |

**<u>Supplementary Table 2:</u>** *ANOVA* output results of the linear regression models used to test the influence of compartments, experimental design and substrate condition on the fungal richness (***chao1*** estimator). To test the influence of compartment and experimental design we used all samples and implemented the following linear regression: *chao1 ~ compartment * exp. design*, where *compartment* is either bulk soil, rhizosphere, roots, leaves or seeds and *exp. design* is either *in situ* or *in vitro*. To test the influence of compartment and substrate condition on *in vitro* seedlings we used the following linear regression: *chao1 ~ compartment * subs. condition*, where *compartment* is either roots or leaves and *subs. condition* is either autoclaved or non-autoclaved. We then used Tukey post-hoc multiple comparisons test for pairwise comparisons.

| **All samples** | **Df** | **Sum Sq** | **Mean Sq** | **F value** | **Pr(>F)** |
|---|---|---|---|---|---|
| compartment | 4 | 27873 | 6968 | 28.083 | **<2,00E-16** |
| exp. design | 1 | 3253 | 3253 | 13.110 | **0.000355** |
| compartment:exp. design | 1 | 934 | 934 | 3.764 | 0.053492 |
| Residuals | 252 | 62529 | 248 | | |
| ***In vitro*** | **Df** | **Sum Sq** | **Mean Sq** | **F value** | **Pr(>F)** |
| compartment | 1 | 87 | 87.1 | 0.393 | 0.535 |
| substrate condition | 1 | 371 | 371.0 | 1.674 | 0.204 |
| compartment: subs.condition | 1 | 17 | 17.3 | 0.078 | 0.782 |
| Residuals | 35 | 7758 | 221.7 | | |

**<u>Supplementary Table 3:</u>** *ANOVA* output results of the linear regression models used to test the influence of compartments, experimental design and substrate condition on the fungal diversity (***Shannon*** index). To test the influence of compartment and experimental design we used all samples and implemented the following linear regression: *Shannon ~ compartment * exp. design*, where *compartment* is either bulk soil, rhizosphere, roots, leaves or seeds and *exp. design* is either *in situ* or *in vitro*. To test the influence of compartment and substrate condition on *in vitro* seedlings we used the following linear regression: *Shannon ~ compartment * subs. condition*, where *compartment* is either roots or leaves and *subs. condition* is either autoclaved or non-autoclaved. We then used Tukey post-hoc multiple comparisons test for pairwise comparisons.

| **All samples** | **Df** | **Sum Sq** | **Mean Sq** | **F value** | **Pr(>F)** |
|---|---|---|---|---|---|
| compartment | 4 | 19.66 | 4.915 | 10.544 | **6.48e-08** |
| exp. design | 1 | 1.43 | 1.428 | 3.063 | 0.0813 |
| compartment:exp. design | 1 | 2.14 | 2.141 | 4.593 | **0.0331** |
| Residuals | 252 | 117.47 | 0.466 | | |
| ***In vitro*** | **Df** | **Sum Sq** | **Mean Sq** | **F value** | **Pr(>F)** |
| compartment | 1 | 0.216 | 0.2163 | 0.252 | 0.619 |
| subs. condition | 1 | 0.339 | 0.3388 | 0.394 | 0.534 |
| compartment: subs.condition | 1 | 0.000 | 0.0001 | 0.000 | 0.994 |
| Residuals | 35 | 30.081 | 0.8594 | | |

**<u>Supplementary Table 4:</u>** Mean contribution of bulk soil, seeds and the unknown source to the mycobiota estimated using the source tracking algorithm *FEAST* (Shenhav *et al.*, 2019). Numbers in the table are given as percentages (%).

| **Sampling** | **Compartment** | **Condition** | **Bulk soil** | **Seeds** | **Unknown** |
|---|---|---|---|---|---|
| *In situ* | Rhizosphere | | 39 | 2.0 | 59 |
| | Roots | | 32 | 0.82 | 67 |
| | Leaves | | 12 | 48 | 40 |
| *In vitro* | Roots | Autoclaved | 3.2 | 26 | 71 |
| | | N-Autoclaved | 0.92 | 16 | 83 |
| | Leaves | Autoclaved | 4.0 | 33 | 63 |
| | | N-Autoclaved | 0.09 | 17 | 83 |

**Supplementary Table 5:** *ANOVA* output results for the *FEAST* source tracking analysis (Shenhav *et al.*, 2019). We ran the regression with (i) only *in-situ* samples: ORQ(*proportion*) ~ *Source* * *compartment*, where *Source* is either bulk soil, seeds or an unknown source and (ii) with germination samples, seeds and bulk soils used for the *in-vitro* experiment: ORQ(*proportion*) ~ *Source* * *compartment* * *condition*. ORQ is the ordered quantile normalization (Peterson and Cavanaugh, 2020). Diagnostic plots are in Supp. Fig. 6b & 6c.

| ***In situ*** | **Df** | **Sum Sq** | **Mean Sq** | **F value** | **Pr(>F)** |
|---|---|---|---|---|---|
| Source | 2 | 166.7 | 83.35 | 209.676 | **< 2,00E-16** |
| compartment | 2 | 3.7 | 1.85 | 4.658 | **0.00987** |
| Source:compartment | 4 | 148.9 | 37.23 | 93.641 | **< 2,00E-16** |
| Residuals | 540 | 214.7 | 0.40 | | |
| ***In vitro*** | **Df** | **Sum Sq** | **Mean Sq** | **F value** | **Pr(>F)** |
| Source | 2 | 75.94 | 37.97 | 127.587 | **<2e-16** |
| compartment | 1 | 0.06 | 0.06 | 0.199 | 0.6569 |
| substrate condition | 1 | 0.01 | 0.01 | 0.022 | 0.8824 |
| Source:compartment | 2 | 0.31 | 0.16 | 0.523 | 0.5944 |
| Source:subs. condition | 2 | 1.61 | 0.80 | 2.702 | 0.0717 |
| compartment: subs.condition | 1 | 0.00 | 0.00 | 0.009 | 0.9257 |
| Source:compartment:subs.condition | 2 | 0.65 | 0.32 | 1.086 | 0.3412 |
| Residuals | 105 | 31.25 | 0.30 | | |

**Supplementary Table 6:** Number of OTUs shared between sources and sinks (left) and their mean share in the compartment’s mycobiota (right). We defined shared OTUs as OTUs shared between a source and sink (e.g., bulk soil and roots) but excluding the other source (seeds in this example).

| Sampling | Compartment | Condition | Bulk soil | Seeds |
|---|---|---|---|---|
| *In situ* | Rhizosphere | | 229 / 43% | 9 / 0.7% |
| | Roots | | 159 / 40% | 6 / 1.5% |
| | Leaves | | 46 / 6.8% | 54 / 7.9% |
| *In vitro* | Roots | Autoclaved | 4 / <0.01% | 32 / 22.3% |
| | | N-Autoclaved | 0 / 0% | 59 / 22.3% |
| | Leaves | Autoclaved | 1 / <0.01% | 22 / 22.1% |
| | | N-Autoclaved | 0 / 0% | 29 / 22.3% |

#### Supplementary Figures (1- 8)

**<u>Supplementary Figure 1:</u> Histogram of the samples sequencing depths.**

The x-axis corresponds to the total number of reads per sample and the y-axis to the number of samples for each class of sequencing depth.

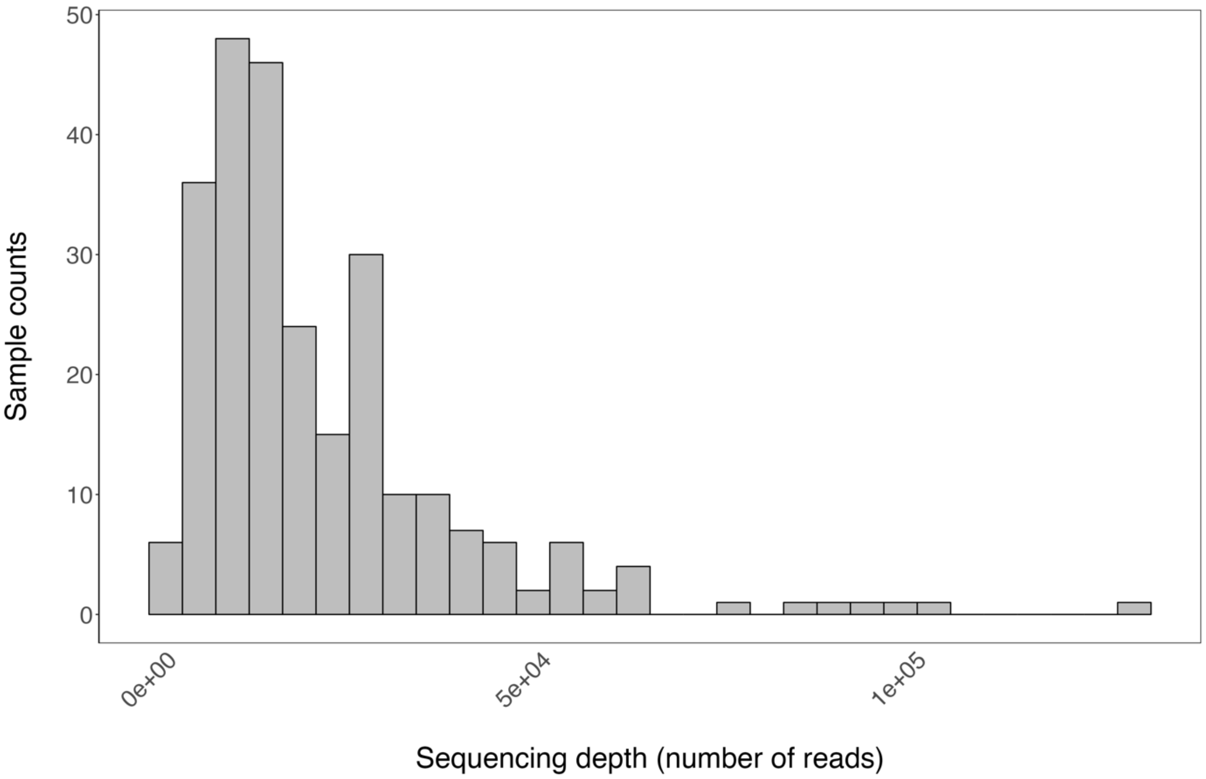

**<u>Supplementary Figure 2:</u> Rarefaction curves for each sample, in each experimental design, for each compartment and each substrate condition *in vitro*.**

The x-axis represents the number of reads and the y-axis the expected richness (number of OTUs). Rarefaction curves were computed using the *rarefy* function of the *vegan* R package (Oksanen *et al.*, 2013).

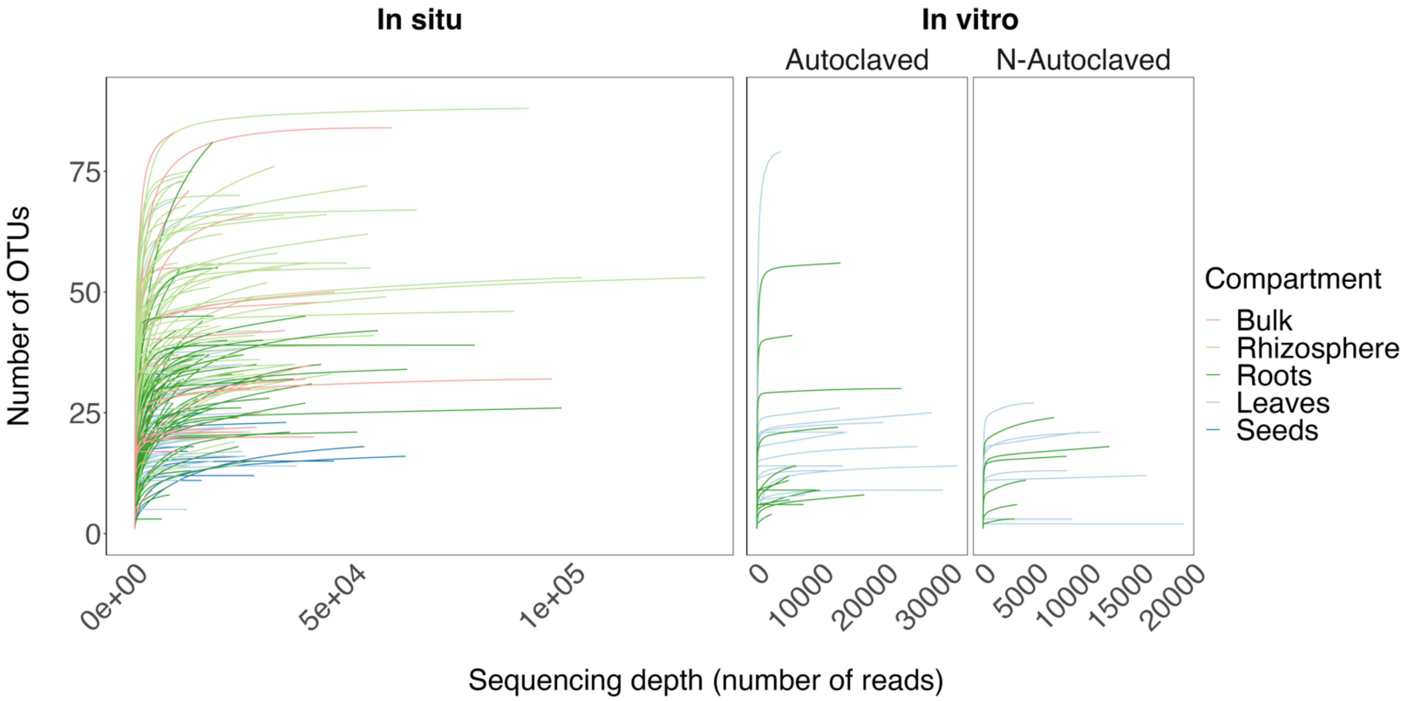

**<u>Supplementary Figure 3:</u> Fungal diversity (Shannon index) of bulk soil and *H. salicornicum* compartments in each experimental design and substrate condition *in vitro*.**

Fungal diversity (*Shannon* index) in the different compartments, experimental designs (*in situ* and *in vitro*) and substrate conditions *in vitro*. Different letters indicate significant differences ($p<0.05$; post-hoc Tukey's pairwise test). Sources (bulk soil and seeds) are in bold font.

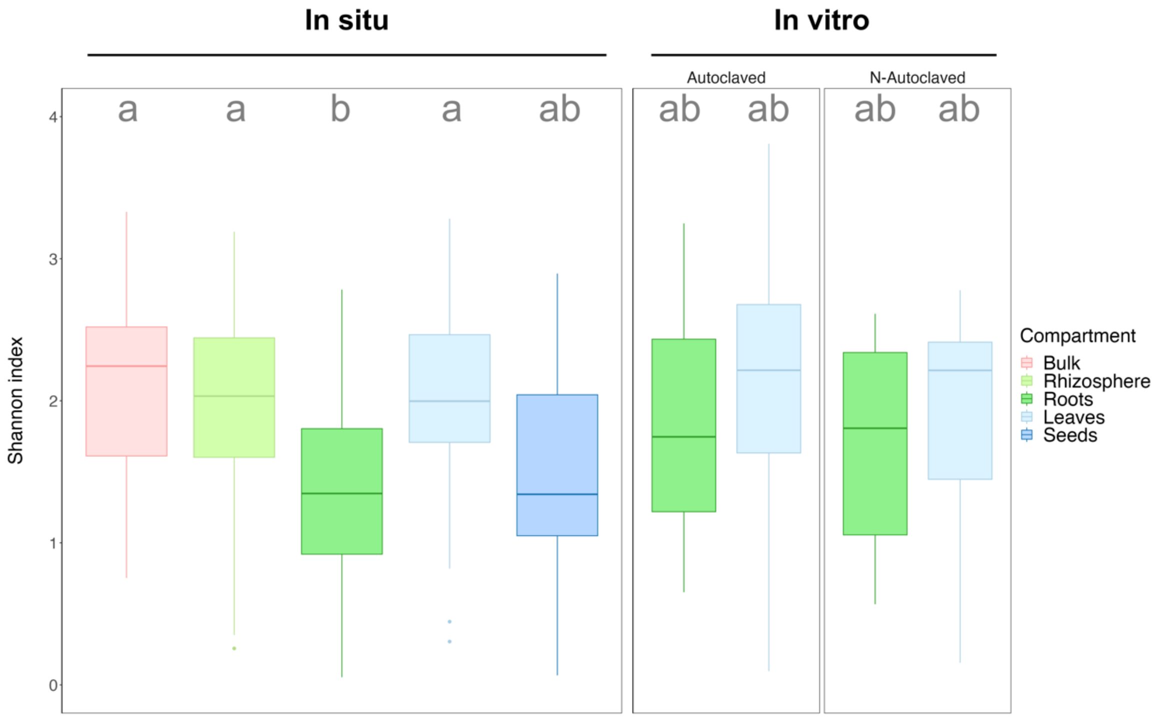

**<u>Supplementary Figure 4:</u> Differences in mycobiota composition between experimental designs (*in situ* and *in vitro*), compartments and substrate conditions (for the *in vitro* experiment) are similar when using UniFrac distances instead of Bray-Curtis distances and when using Hellinger-transformed data instead of Relative Abundances.**

Principal Coordinates Analysis (PCoA) based on Bray-Curtis or UniFrac distances between samples with relative abundances or Hellinger-transformed abundances. Ellipses represent the normal-probability contours of the data. Influence of the variables on distance matrices were tested using *PERMANOVA* (10 000 permutations). The procedure was run with all samples (Total dataset; left) and only for the *in vitro* samples (right).

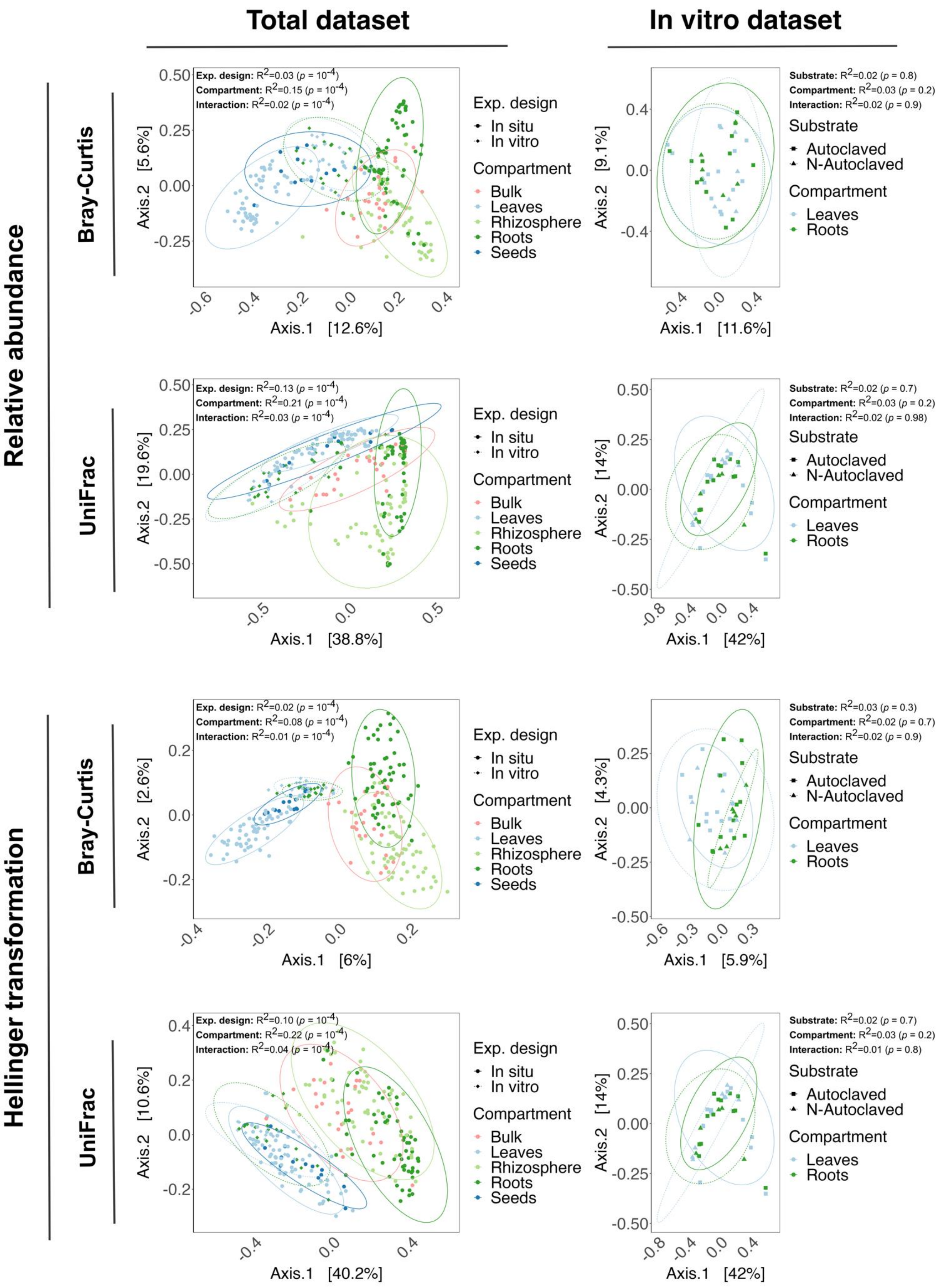
Total dataset
In vitro dataset
Relative abundance
Hellinger transformation
Bray-Curtis
UniFrac
Exp. design: R2=0.03 (p = 10-4)
Compartment: R2=0.15 (p = 10-4)
Interaction: R2=0.02 (p = 10-4)
Exp. design
In situ
In vitro
Compartment
Bulk
Leaves
Rhizosphere
Roots
Seeds
Axis.1 [12.6%]
Axis.2 [5.6%]
Substrate: R2=0.02 (p = 0.8)
Compartment: R2=0.03 (p = 0.2)
Interaction: R2=0.02 (p = 0.9)
Substrate
Autoclaved
N-Autoclaved
Axis.1 [11.6%]
Axis.2 [9.1%]
Exp. design: R2=0.13 (p = 10-4)
Compartment: R2=0.21 (p = 10-4)
Interaction: R2=0.03 (p = 10-4)
Axis.1 [38.8%]
Axis.2 [19.6%]
Substrate: R2=0.02 (p = 0.7)
Compartment: R2=0.03 (p = 0.2)
Interaction: R2=0.02 (p = 0.98)
Axis.1 [42%]
Axis.2 [14%]
Exp. design: R2=0.02 (p = 10-4)
Compartment: R2=0.08 (p = 10-4)
Interaction: R2=0.01 (p = 10-4)
Axis.1 [6%]
Axis.2 [2.6%]
Substrate: R2=0.03 (p = 0.3)
Compartment: R2=0.02 (p = 0.7)
Interaction: R2=0.02 (p = 0.9)
Axis.1 [5.9%]
Axis.2 [4.3%]
Exp. design: R2=0.10 (p = 10-4)
Compartment: R2=0.22 (p = 10-4)
Interaction: R2=0.04 (p = 10-4)
Axis.1 [40.2%]
Axis.2 [10.6%]
Substrate: R2=0.02 (p = 0.7)
Compartment: R2=0.03 (p = 0.2)
Interaction: R2=0.01 (p = 0.8)
Axis.1 [42%]
Axis.2 [14%]

__Supplementary Figure 5:__ **Bipartite networks for each compartment in each dataset and condition.** Grey nodes represent OTUs and colored ones samples. Diameter of the nodes is proportional to the betweenness centrality. Width of the edges is proportional to the relative abundance of an OTU in a sample. We used the Fruchterman–Reingold layout algorithm for better readability (Fruchterman and Reingold, 1991). See Supp. Methods 3 for metrics calculation and their significance tests. **(a)** *In situ* samples. **(b)** *In vitro* samples.

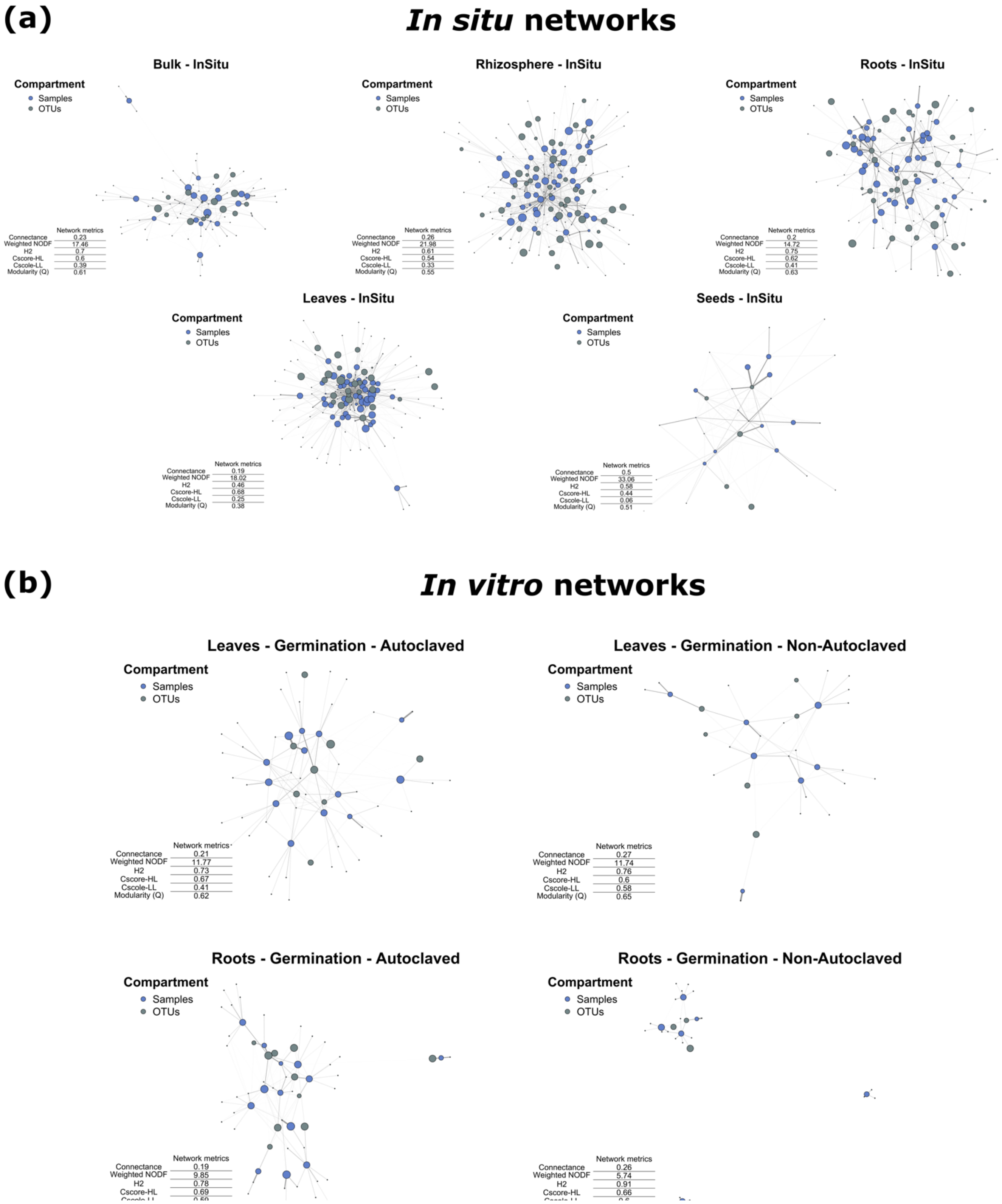

(a)
In situ networks
Bulk - InSitu
Rhizosphere - InSitu
Roots - InSitu
Leaves - InSitu
Seeds - InSitu
Compartment
Samples
OTUs
Network metrics
Connectance 0.23
Weighted NODF 17.46
H2 0.7
Cscore-HL 0.6
Cscole-LL 0.39
Modularity (Q) 0.61
Connectance 0.26
Weighted NODF 21.98
H2 0.61
Cscore-HL 0.54
Cscole-LL 0.33
Modularity (Q) 0.55
Connectance 0.2
Weighted NODF 14.72
H2 0.75
Cscore-HL 0.62
Cscole-LL 0.41
Modularity (Q) 0.63
Connectance 0.19
Weighted NODF 18.02
H2 0.46
Cscore-HL 0.68
Cscole-LL 0.25
Modularity (Q) 0.38
Connectance 0.5
Weighted NODF 33.06
H2 0.58
Cscore-HL 0.44
Cscole-LL 0.06
Modularity (Q) 0.51
(b)
In vitro networks
Leaves - Germination - Autoclaved
Leaves - Germination - Non-Autoclaved
Roots - Germination - Autoclaved
Roots - Germination - Non-Autoclaved
Connectance 0.21
Weighted NODF 11.77
H2 0.73
Cscore-HL 0.67
Cscole-LL 0.41
Modularity (Q) 0.62
Connectance 0.27
Weighted NODF 11.74
H2 0.76
Cscore-HL 0.6
Cscole-LL 0.58
Modularity (Q) 0.65
Connectance 0.19
Weighted NODF 9.85
H2 0.78
Cscore-HL 0.69
Connectance 0.26
Weighted NODF 5.74
H2 0.91
Cscore-HL 0.66

**<u>Supplementary Figure 6:</u> Transformation of *FEAST* outputs used to test the influence of source, compartment and condition on the contribution to the mycobiota.**
**(a)** Ordered quantile normalization (ORQ; Peterson and Cavanaugh, 2020) of the proportions of the mycobiota explained by the potential sources (bulk soil, seeds and an unknow source). **(b), (c)** Diagnostic plots for the two regressions used. **(b)** Regression with only samples *in situ*: ORQ(*proportion*) ~ *source* * *compartment*, where *source* is either bulk soil, seeds or an unknown source. **(c)** Regression with samples *in vitro*, seeds and bulk soils used for the *in-vitro* experiment: ORQ(*proportion*) ~ *source* * *compartment* * *condition*.

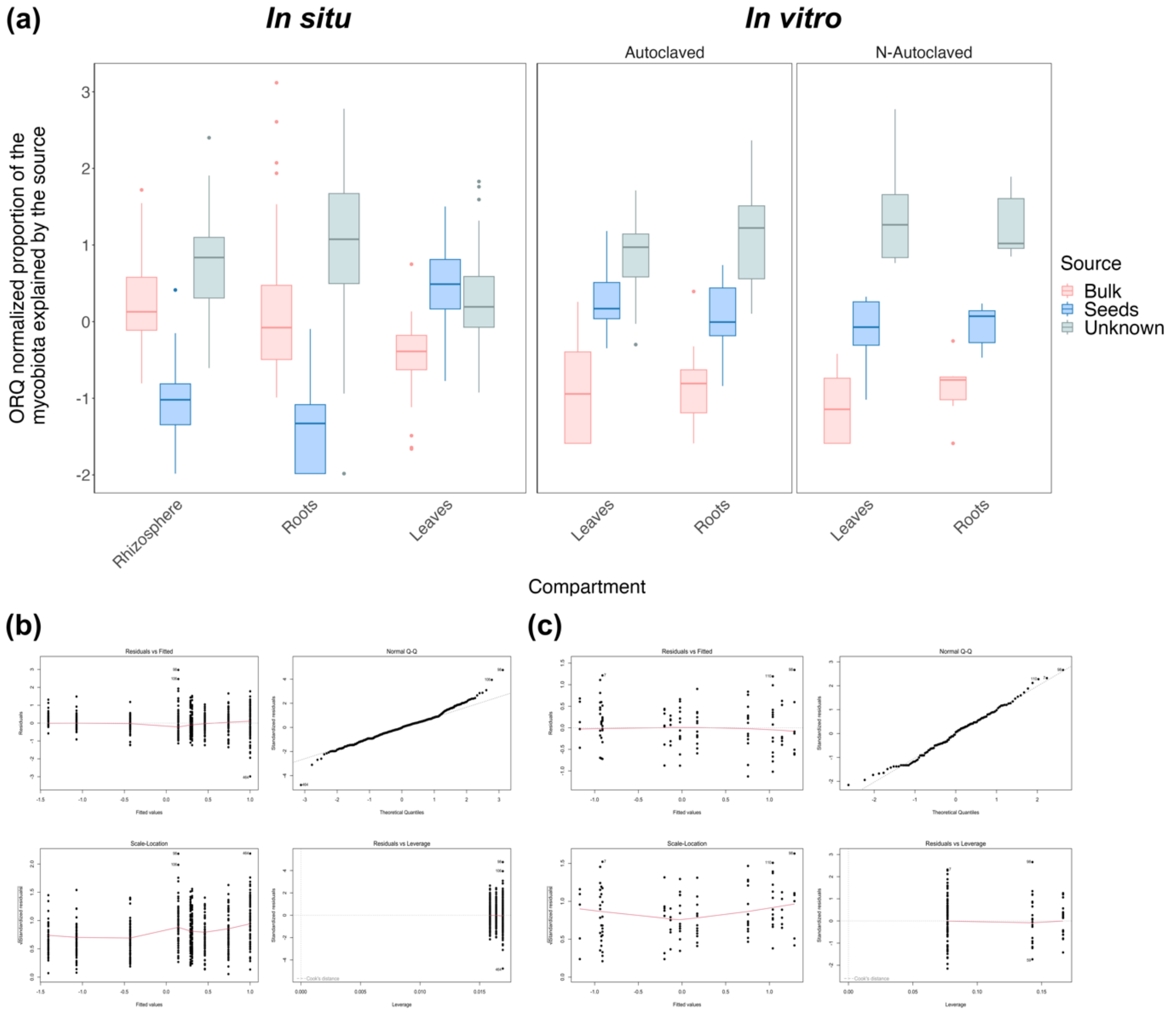

**<u>Supplementary Figure 7:</u> Potentially transmitted fungi by seeds and bulk soil when considering ubiquitous fungi.**

Mycobiota composition at the Genus level of each compartment in each dataset and condition when only considering OTUs shared between the sink (rhizosphere, roots or leaves) and the potential source (bulk or seeds) without excluding OTUs shared between the two sources (i.e., we only take into consideration OTUs shared between the studied compartment and the potential source). These are the potentially transmitted OTUs from sources to sinks.

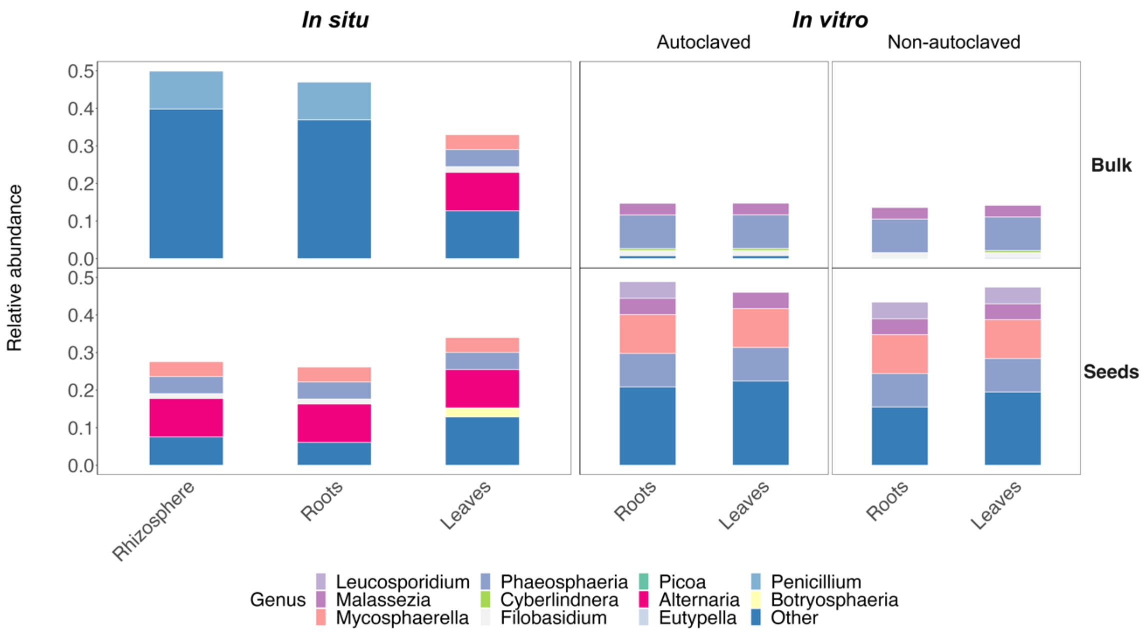

**<u>Supplementary Figure 8:</u> Both the proportion of the mycobiota explained by a source and to the number of shared OTUs between this source & a given compartment are positively correlated to the number of shared OTUs between the source & the compartment studied.** The explained proportion of the mycobiota is the estimated proportion using the source-tracking algorithm *FEAST* (Shenhav *et al.*, 2019). We defined shared OTUs as OTUs shared between a source and a sink (e.g., bulk soil and roots) but excluding the other source (seeds in this example). The mean proportion of the shared mycobiota correspond to the mean values of the total relative abundance represented by the shared OTUs (as defined before) in the compartment of interest (e.g., the mean proportion of root mycobiota when considering shared OTUs between roots and bulk soil). **(a)** Linear regression between the number of shared OTUs (log positive transformed) and the estimated proportion of the mycobiota explained by the sources (log positive transformed). **(b)** Linear regression between the mean proportion of the shared OTUs and the estimated proportion of the mycobiota explained by the sources.

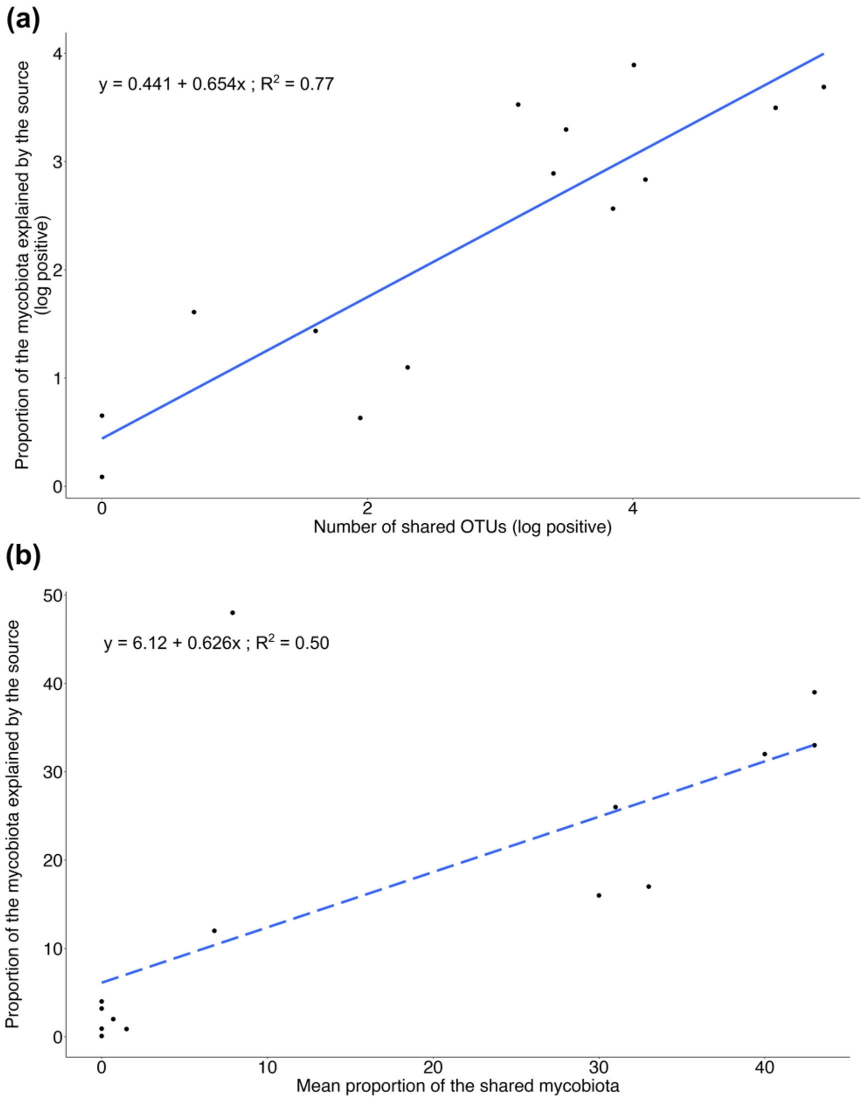

#### References

Blüthgen, Nico, Menzel, F., and Blüthgen, Nils (2006) Measuring specialization in species interaction networks. *BMC Ecol* **6**: 9.

Capella-Gutiérrez, S., Silla-Martínez, J.M., and Gabaldón, T. (2009) trimAl: a tool for automated alignment trimming in large-scale phylogenetic analyses. *Bioinformatics* **25**: 1972–1973.

Csardi, G. and Nepusz, T. (2006) The igraph software package for complex network research. *InterJournal Complex Syst* **1695**: 1–9.

Dormann, C.F., Gruber, B., and Fründ, J. (2008) Introducing the bipartite package: analysing ecological networks. *interaction* **1**: 8–11.

Fruchterman, T.M.J. and Reingold, E.M. (1991) Graph drawing by force-directed placement. *Softw Pract Exp* **21**: 1129–1164.

Katoh, K. and Standley, D.M. (2013) MAFFT Multiple Sequence Alignment Software Version 7: Improvements in Performance and Usability. *Mol Biol Evol* **30**: 772–780.

Martin, M. (2011) Cutadapt removes adapter sequences from high-throughput sequencing reads. *EMBnet.journal* **17**: 10.

Maurice, K., Bourceret, A., Youssef, S., Boivin, S., Laurent-Webb, L., Damasion, C., et al. (2023) Anthropic disturbances impact the soil microbial network structure and stability to a greater extent than natural disturbances in an arid ecosystem. *Manuscript submitted for publication*.

McMurdie, P.J. and Holmes, S. (2013) phyloseq: An R Package for Reproducible Interactive Analysis and Graphics of Microbiome Census Data. *PLOS ONE* **8**: e61217.

Nguyen, L.-T., Schmidt, H.A., von Haeseler, A., and Minh, B.Q. (2015) IQ-TREE: A Fast and Effective Stochastic Algorithm for Estimating Maximum-Likelihood Phylogenies. *Mol Biol Evol* **32**: 268–274.

Nilsson, R.H., Larsson, K.-H., Taylor, A.F.S., Bengtsson-Palme, J., Jeppesen, T.S., Schigel, D., et al. (2019) The UNITE database for molecular identification of fungi: handling dark taxa and parallel taxonomic classifications. *Nucleic Acids Res* **47**: D259–D264.

Oksanen, J., Blanchet, F.G., Kindt, R., Legendre, P., Minchin, P.R., O'hara, R.B., et al. (2013) Package 'vegan.' *Community Ecol Package Version* **2**: 1–295.

Op De Beeck, M., Lievens, B., Busschaert, P., Declerck, S., Vangronsveld, J., and Colpaert, J.V. (2014) Comparison and Validation of Some ITS Primer Pairs Useful for Fungal Metabarcoding Studies. *PLoS ONE* **9**: e97629.

Perez-Lamarque, B., Krehenwinkel, H., Gillespie, R.G., and Morlon, H. (2022) Limited Evidence for Microbial Transmission in the Phylosymbiosis between Hawaiian Spiders and Their Microbiota. *mSystems* **7**: e01104-21.

Perez-Lamarque, B., Laurent-Webb, L., Bourceret, A., Maillet, L., Bik, F., Cartier, D., et al. (2023) Fungal microbiomes associated with Lycopodiaceae during ecological succession. *Environ Microbiol Rep* **15**: 109–118.

Perez-Lamarque, B., Petrolli, R., Strullu-Derrien, C., Strasberg, D., Morlon, H., Selosse, M.-A., and Martos, F. (2022) Structure and specialization of mycorrhizal networks in phylogenetically diverse tropical communities. *Environ Microbiome* **17**: 38.

Peterson, R.A. and Cavanaugh, J.E. (2020) Ordered quantile normalization: a semiparametric transformation built for the cross-validation era. *J Appl Stat* **47**: 2312–2327.

Petrolli, R., Augusto Vieira, C., Jakalski, M., Bocayuva, M.F., Vallé, C., Cruz, E.D.S., et al. (2021) A fine-scale spatial analysis of fungal communities on tropical tree bark unveils the epiphytic rhizosphere in orchids. *New Phytol* **231**: 2002–2014.

R Core Team (2023) R: A Language and Environment for Statistical Computing, Vienna, Austria: R Foundation for Statistical Computing.

Rognes, T., Flouri, T., Nichols, B., Quince, C., and Mahé, F. (2016) VSEARCH: a versatile open source tool for metagenomics. *PeerJ* **4**: e2584.

Shenhav, L., Thompson, M., Joseph, T.A., Briscoe, L., Furman, O., Bogumil, D., et al. (2019) FEAST: fast expectation-maximization for microbial source tracking. *Nat Methods* **16**: 627–632.

Susana Rivera, C., Eugenia Venturini, M., Oria, R., and Blanco, D. (2011) Selection of a decontamination treatment for fresh Tuber aestivum and Tuber melanosporum truffles packaged in modified atmospheres. *FOOD CONTROL* **22**: 626–632.

Tedersoo, L., Bahram, M., Zinger, L., Nilsson, R.H., Kennedy, P.G., Yang, T., et al. (2022) Best practices in metabarcoding of fungi: From experimental design to results. *Mol Ecol* **31**: 2769–2795.

White, T.J., Bruns, T., Lee, S., and Taylor, J. (1990) AMPLIFICATION AND DIRECT SEQUENCING OF FUNGAL RIBOSOMAL RNA GENES FOR PHYLOGENETICS. In *PCR Protocols*. Elsevier, pp. 315–322.
